# A directed TM→JM coupling in receptor tyrosine kinase dimers, set by activating mutations and the membrane environment

**DOI:** 10.64898/2026.05.22.727318

**Authors:** Takeshi Sato, Hiroko Tamagaki-Asahina

**Affiliations:** Kyoto Pharmaceutical University, Kyoto 607-8414, Japan

## Abstract

The direction of conformational coupling in a membrane protein, that is, which domain drives which, has been inaccessible to experiment. We recover this directivity from molecular dynamics (MD) of transmembrane–juxtamembrane (TM–JM) dimers of receptor tyrosine kinases EGFR and FGFR3. Coupling is detected with a Bayesian-network framework (CASCADE); its direction is measured with PERI (Phase-plane Estimation of Rotational Irreversibility), the net phase-plane circulation, validated on synthetic data and resolved at 0.1 ns. Direction is summarized as the TM→JM directed-mass fraction f+ (0.5 = balanced) via a hierarchical Bayesian model. The activating TM mutants EGFR L658Q and FGFR3 A391E are TM→JM (posterior probability 0.95 and 0.99); fluid wild-type EGFR leans the same way (0.93), in agreement with its experimentally reported constitutive activity in fluid but not ordered bilayers; the ligand-dependent ordered wild type is balanced (0.45); and an activating mutation raises the TM→JM bias above the ordered wild type with probability 0.94. The directivity thus tracks the measured activity state of the receptor, distinguishing signaling-competent from ligand-dependent RTK dimers by a property not apparent from structure alone.

## Introduction

Receptor tyrosine kinases (RTKs) transduce extracellular signals through dimerization and activation of their intracellular kinase domains.^1, 2^ Membrane lipid composition profoundly influences this process: lipid rafts modulate FGFR2 signaling,^3^ the ganglioside GM3 regulates the insulin receptor^4^ and inhibits EGFR dimerization.^5^ Most directly, in reconstituted membranes the EGF receptor is constitutively active in fluid bilayers but requires ligand for activation in ordered, raft-like bilayers.^5^ The membrane thus controls RTK activity, but the conformational mechanism through which it does so is not known.

Within an RTK dimer the transmembrane (TM) helix geometry and the positioning of the intracellular juxtamembrane (JM) segment are conformationally coupled^6-9^. Structural biology has long assumed a direction for this coupling: ligand-induced rotation of the TM domain propagates outward-to-inward to the JM segment and releases it from the membrane to activate the kinase.^10^ Whether the structural fluctuations within a single dimer actually carry a measurable direction, whether that direction can change, and whether the membrane controls it, have never been asked, because the direction is experimentally inaccessible. Ensemble spectroscopies such as solid-state NMR resolve domain orientation and inter-domain coupling but average over all molecules in the sample; single-molecule fluorescence can resolve dynamics but cannot place time-resolved probes on both domains of both chains within one dimer without a labeling density incompatible with a single-pass peptide. Molecular dynamics (MD) provides the required per-frame, per-variable resolution,^11, 12^ yet the analyses usually applied to trajectories, such as principal component analysis,^13, 14^ mutual information,^15, 16^ and dynamic network analysis^17^, quantify how strongly variables co-fluctuate without resolving which one leads. If two membrane conditions produce the same covariation between the same structural variables but differ in which variable drives which, no such method can tell them apart.

This work builds on two decades of experimental structural studies of RTK TM–JM peptides by solid-state NMR, ATR-IR, and fluorescence spectroscopy,^7-9^ which characterized the TM–JM coupling but could not resolve its direction of propagation. We separate the problem into two questions. First, do a pair of structural variables co-fluctuate at all: is there coupling to which a direction can be assigned? We answer this with CASCADE, a Bayesian-network structural-covariance framework used here only for coupling detection^18-20^ that returns the undirected coupling skeleton among structural variables and hierarchically removes the dominant membrane-order signal so that internal protein coupling becomes visible (its edge orientations are not used; direction is assigned by PERI). Second, given coupling, which domain leads? Rather than orient observational graph edges, which is in part a modeling choice^18^, we measure the direction with PERI (Phase-plane Estimation of Rotational Irreversibility): the net phase-plane circulation of the two coupled domains, the net probability current, which is the physically correct, time-reversal-odd signature of a directed (irreversible) process (Figure 1). We validate PERI on synthetic data with known direction and then apply it to TM–JM dimers of EGFR and FGFR3, chosen because high-resolution NMR structures of their TM–JM segments have been determined^6, 21^, both harbor clinically significant disease-associated TM mutations,^22, 23^ and their TM domains share less than 20% sequence identity.

**Figure 1.**
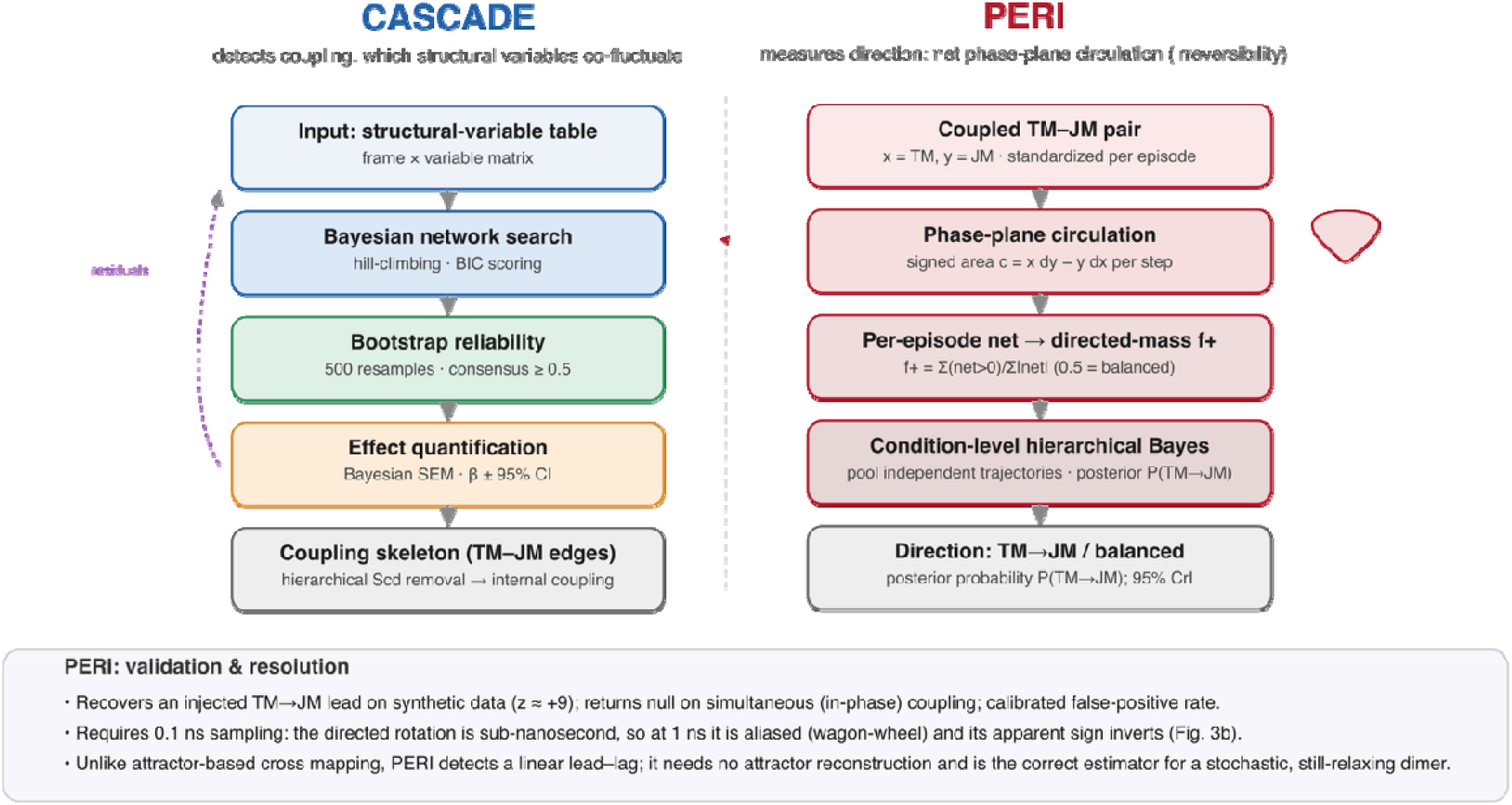
The two-stage framework. CASCADE detects coupling (which structural variables co-fluctuate) by Bayesian-network search with bootstrap reliability and hierarchical removal of the membrane-order signal. PERI (Phase-plane Estimation of Rotational Irreversibility) then measures direction as the net phase-plane circulation of a coupled TM–JM pair (signed area c = x dy − y dx; net probability current), summarized by the directed-mass fraction f+ and inferred at the condition level with a hierarchical Bayesian model. PERI recovers an injected TM→JM lead on synthetic data, is null on simultaneous coupling, and is applied at 0.1 ns, which resolves the fast (sub-nanosecond) TM→JM component of the scale-dependent coupling.

## Results

### Detecting coupling, then measuring its direction

CASCADE takes structural variables measured at each frame and identifies which co-fluctuate, using hill-climbing search with Bayesian information scoring, bootstrap edge stability, and Bayesian structural-equation effect sizes; a hierarchical mode removes a dominant environmental variable (local membrane order, the lipid acyl-chain order parameter Scd) and reruns on the residuals to expose internal protein coupling. CASCADE thus provides the coupling skeleton, the pairs of TM and JM variables that move together, but not a reliable direction, consistent with the known limitation that observational edge orientation is partly a modeling choice^18, 19^.

Direction is assigned separately, by PERI (Phase-plane Estimation of Rotational Irreversibility; Figure 1). For a TM variable x and a JM variable y, standardized within each association episode (a contiguous period during which the two protomers remain in TM contact; defined in Methods), we compute the signed area swept per step in the (x, y) plane, c(t) = x_mΔy − y_mΔx, and sum it: net > 0 means the joint state circulates such that TM leads JM (TM→JM); net < 0 means JM→TM; a simultaneous, in-phase coupling encloses no area and gives net ≈ 0. This net circulation is the net probability current of the two-domain dynamics, nonzero only when detailed balance is broken, and it changes sign under time reversal, the defining property of a directed process; this phase-plane circulation is the established signature of broken detailed balance in stochastic systems.^24^ Against a within-episode circular-shift null (which preserves the distribution and autocorrelation of each variable while destroying only the TM–JM alignment), the statistic recovers an injected TM→JM lead (z ≈ +9.8), returns null on simultaneous coupling, and holds its false-positive rate close to the nominal level (0.075; SI Table S2) even when the two variables differ in smoothness or share a common slow driver.

### CASCADE resolves the TM–JM coupling skeleton

CASCADE returned a dense but structured coupling network in both membranes (Figure 2), computed from nine structural observables per frame: two local membrane-order parameters (Scd), four TM-geometry variables (the two helix tilts, the crossing angle, and the inter-TM distance) and three intracellular JM-position variables. In the fluid EGFR wild-type dimer the consensus network contained 25 edges and in the ordered dimer 30; in both, the two Scd variables were each coupled to all seven protein variables (bootstrap frequency 0.94–1.00 in fluid, 0.58–1.00 in ordered), identifying local membrane order as the dominant hub (edge bootstrap frequencies, Figure S1; variance explained in each protein variable, Figure S2). Because this environmental variable co-varies with essentially every protein variable, its shared variance was regressed out hierarchically, identically in every condition, and CASCADE was rerun on the residuals to expose the internal protein coupling. The internal network connects TM geometry to JM position through multiple cross-group edges (10 in the fluid wild type, 15 in the ordered wild type), establishing that the TM and JM domains co-fluctuate and providing the skeleton to which a direction can be assigned. In the ordered membrane, on which Figure 2 focuses, the activating L658Q dimer gave a sparser internal network than the ordered wild type (8 edges against 15), consistent with a more constrained, pre-organized dimer.

**Figure 2.**
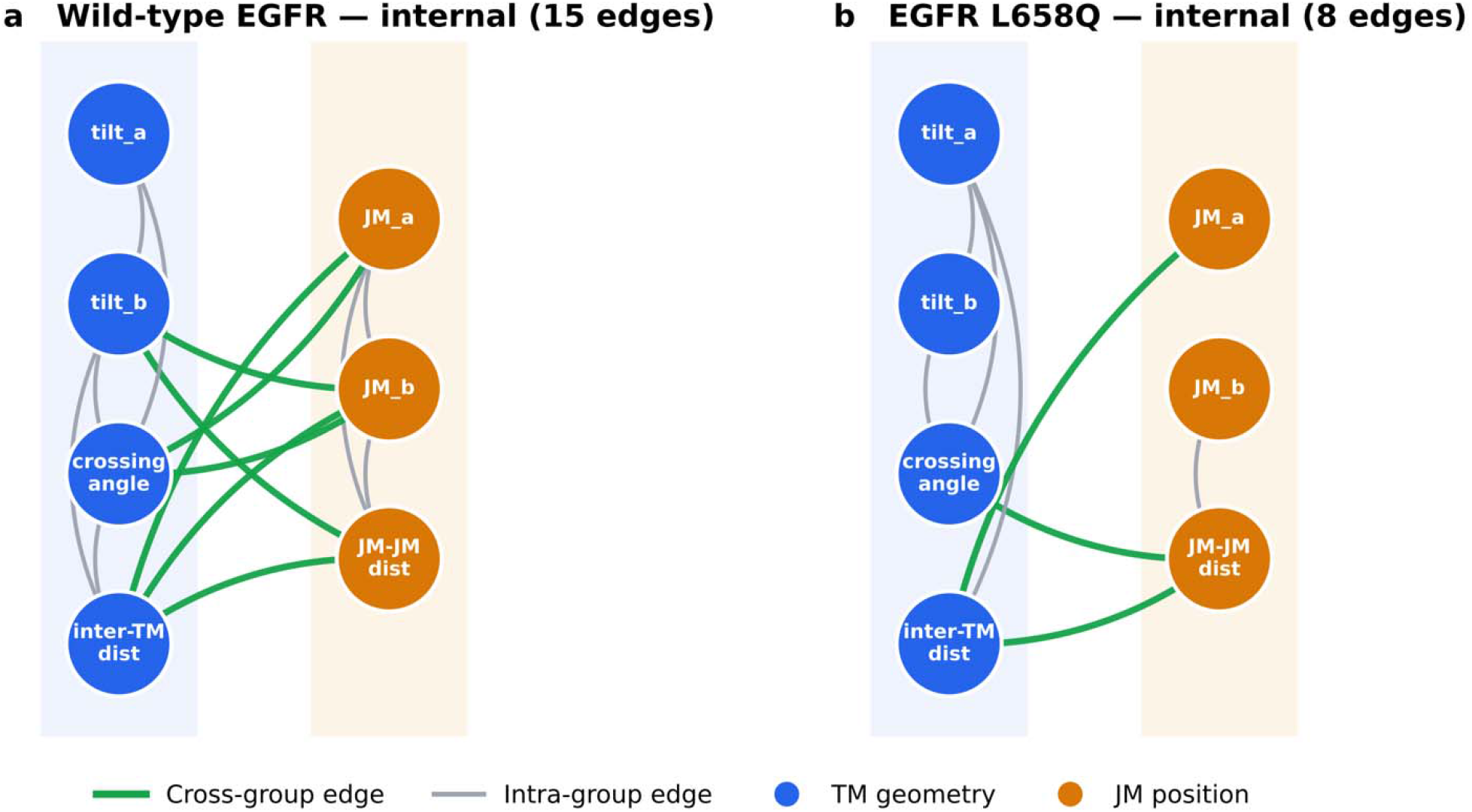
CASCADE coupling networks in the ordered membrane (EGFR in DIPC/DPSM/cholesterol). Internal Bayesian-network skeletons among the structural variables (TM geometry, JM position), shown after hierarchical removal of the dominant membrane-order (Scd) variance to expose the internal TM– JM coupling; edge thickness/appearance reflects bootstrap frequency. (a) Wild-type EGFR retains a dense internal TM–JM network (15 edges); (b) the activating L658Q dimer is sparser (8 edges). Orientation of these edges is not interpreted as direction (Note S1.7); direction is measured by PERI (Figure 3). Full and internal edge counts for both membranes (fluid 25/10, ordered 30/15) are given in the SI.

Intuitively, PERI watches the dimer as a single point moving in a plane whose axes are the TM state and the JM state. If the two domains moved in lockstep the point would slide back and forth along a line and enclose nothing; if one leads the other it traces a loop, and the sense in which the loop turns is the direction of the coupling (one sense is TM→JM, the reverse JM→TM), while the area it encloses is the strength (Figure 3a). Because a dimer emits strong loops of both senses, a condition is read not from any single episode but from the balance between them: the directed-mass fraction f+ is the TM→JM share of the total loop area, so f+ = 0.5 means the two senses cancel (no leader), f+ > 0.5 a net TM→JM lead, and f+ < 0.5 a net JM→TM lead. A few strong, well-formed loops dominate f+ while faint, loop-free episodes count for almost nothing, so f+ reports the direction of the strong association events.

**Figure 3.**
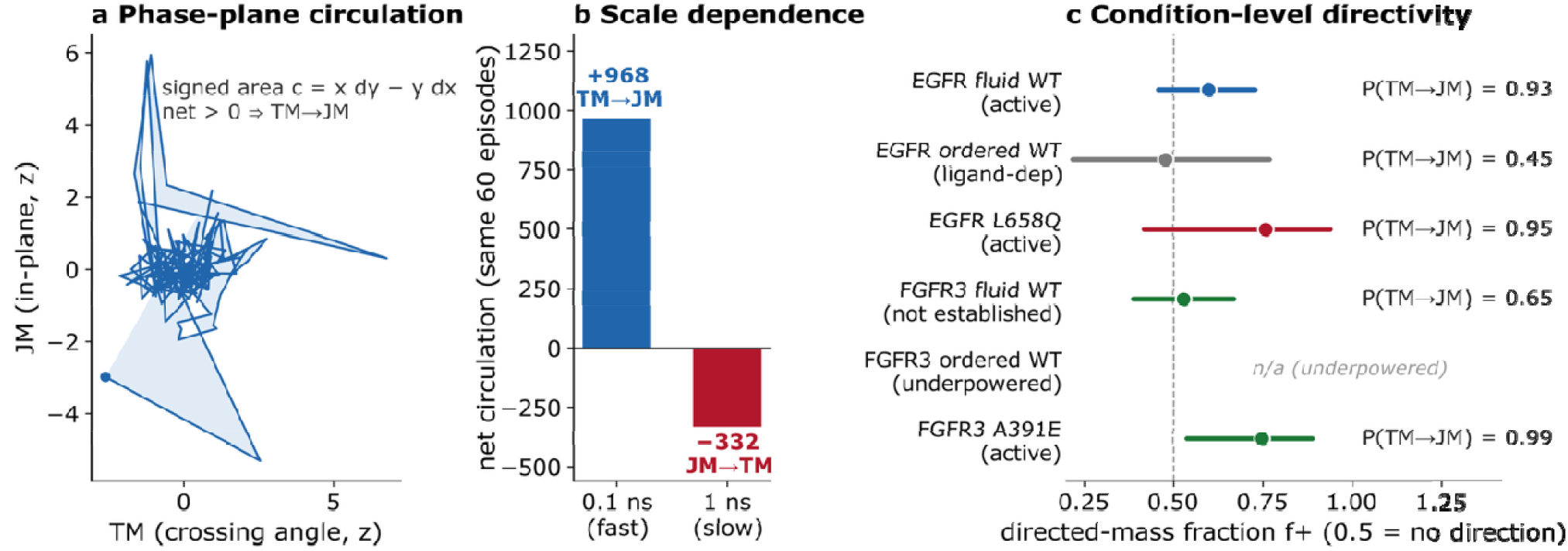
PERI directivity at 0.1 ns. (a) One association episode of the EGFR wild-type fluid dimer traced in the (TM, JM) plane; the enclosed signed area is the net circulation (net > 0 □ TM→JM). (b) On the same 60 episodes of one EGFR fluid trajectory the net circulation is +968 (TM→JM) at 0.1 ns but −332 (JM→TM) at 1 ns: the fast sub-nanosecond TM→JM rotation is aliased at 1 ns and the slow JM→TM rotation dominates, so the apparent sign inverts (scale-dependence: Figure 4). (c) Condition-level directed-mass fraction f+ (0.5 = no direction) with hierarchical Bayesian 95% credible interval; labels give the posterior probability P(TM→JM). The activating TM mutants EGFR L658Q and FGFR3 A391E are TM→JM with posterior probability 0.95 and 0.99, fluid wild-type EGFR 0.93, the ordered wild type balanced (0.45), and wild-type FGFR3 in fluid balanced (0.65); wild-type FGFR3 in the ordered membrane is shown as not available (underpowered; too few discrete episodes).

### The directed coupling is scale-dependent: 0.1 ns resolves the fast component

The directed coupling is scale-dependent, and this dictates the sampling. Decomposing the phase-plane circulation of one wild-type EGFR fluid trajectory by frequency (SI Note S3; Figure 4a) resolves two components of opposite sense: a fast, sub-nanosecond rotation that is TM→JM (band contribution +915) and dominates the full-bandwidth net of +968, and a slower, multi-nanosecond rotation that is JM→TM (band contribution −217); because the phase-plane circulation is a bilinear functional of the two signals, these band contributions include cross terms and do not sum exactly to the full-bandwidth net. The fast component lies above the 1 ns Nyquist limit (0.5 ns□^1^): at 1 ns sampling it is aliased (a stroboscopic “wagon-wheel” effect) and the slow JM→TM rotation dominates the apparent direction, so the same episodes read significantly JM→TM at 1 ns but significantly TM→JM at 0.1 ns (Figure 3b). Both rotations are real; the direction one measures depends on the timescale probed. The fast TM→JM rotation is the direct mechanical coupling between the two covalently connected ends of a single peptide, the intramolecular TM–JM coupling this study measures, and the one aliased at coarser sampling, so all directivity results below use 0.1 ns geometry, at which PERI resolves it (Methods; SI).

**Figure 4.**
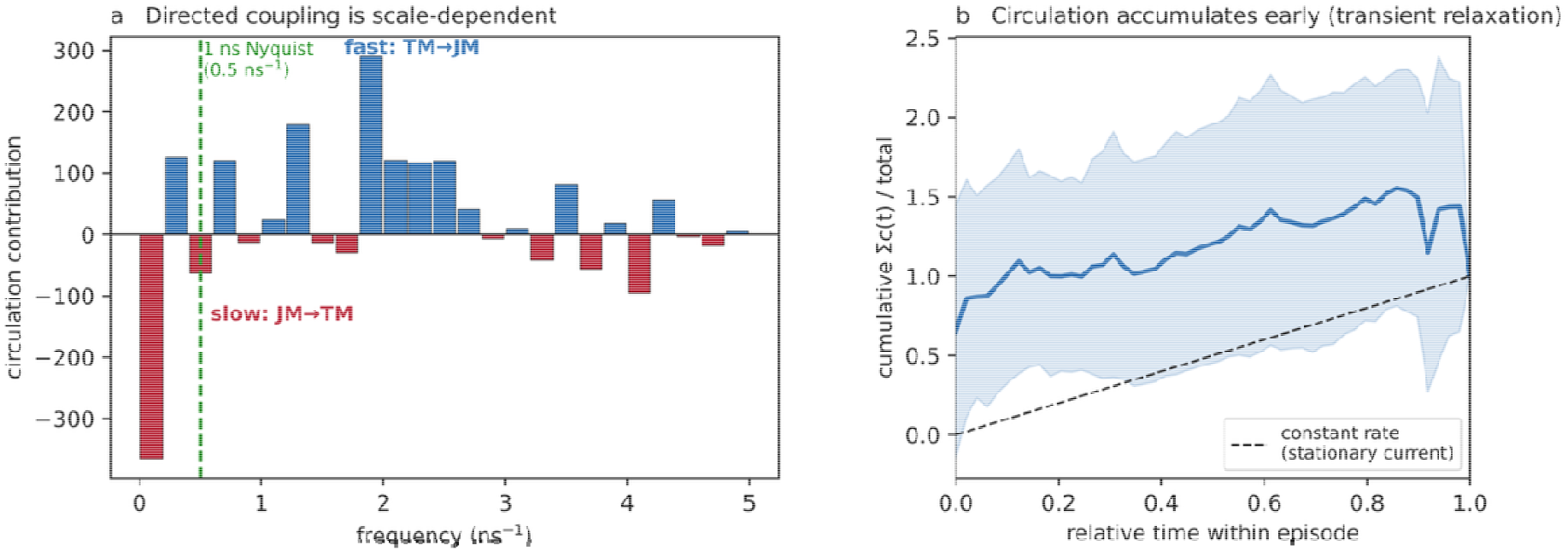
Scale dependence and transient character of the directed circulation (EGFR wild-type fluid, 0.1 ns). (a) Frequency-resolved circulation: below ≈0.5 ns□^1^ (the 1 ns Nyquist limit, dashed) the net is JM→TM (red); above it the net is TM→JM (blue) and carries most of the full-bandwidth signal, so 1 ns sampling aliases the fast TM→JM rotation and reads the slow JM→TM one. (b) Cumulative Σc(t) within an episode (mean ± 95% CI) rises above the constant-rate diagonal, reaching most of its final value early: the directed circulation is a transient of relaxation, not a stationary current.

### TM→JM directivity marks constitutively active EGFR

We performed coarse-grained MD of EGFR TM–JM dimers (residues 634–683, PDB 2M20) in fluid (dilinoleoyl phosphatidylcholine (DIPC) and dioleoyl phosphatidylserine (DOPS)) and ordered (DIPC, sphingomyelin (DPSM), and cholesterol) membranes (GROMACS 2023.3^25^, MARTINI 2.2^26, 27^; no elastic network; see Methods). CASCADE confirmed coupling between TM geometry and intracellular JM position in both membranes (Figure 2). The direction of this coupling is not uniform across association episodes: in every condition the dimer produces strong directed events in both senses (some episodes circulate TM→JM, others JM→TM), while the many low-amplitude episodes contribute essentially no net rotation (they are, as expected, noise and carry ≈0). The directivity of a condition is therefore the net imbalance between the strong TM→JM and strong JM→TM events, summarized by f+ = Σ(positive per-episode net)/Σ|net|, the TM→JM fraction of the directed mass (0.5 = balanced). Because a single trajectory yields few episodes, direction is inferred at the condition level with a hierarchical Bayesian model over the independent trajectories (SI Note S3.4), which returns the posterior probability P(TM→JM) that the condition is TM→JM-biased together with a 95% credible interval (CrI) for f+. At 0.1 ns, wild-type EGFR in fluid membranes, the constitutively active state, is TM→JM-biased in both independent trajectories (f+ = 0.62 and 0.59, same sign), giving f+ = 0.60 (95% CrI 0.46–0.73) and P(TM→JM) = 0.93. The bias toward the kinase is reproducible, consistent with the constitutive activity of EGFR in fluid membranes reported by Coskun et al.^5^

### The ordered membrane balances the two senses in wild-type EGFR

In ordered membranes, where wild-type EGFR requires ligand, the coupling is balanced: the two independent trajectories disagree in sign (f+ = 0.42 and 0.52) and pool to f+ = 0.48 (95% CrI 0.22–0.77; P(TM→JM) = 0.45), a posterior centered on 0.5, the signature of a genuine absence of net direction rather than a weak consistent one. The ordered membrane environment does not invert the coupling; it equalizes the two senses, leaving no net direction, consistent with the ligand dependence of wild-type EGFR in ordered membranes.

### The oncogenic L658Q mutation restores TM→JM directivity

The EGFR L658Q substitution, which causes ligand-independent activation, restored a TM→JM bias in the same ordered membrane that balances it in wild type. Both independent trajectories shift toward TM→JM (f+ = 0.71 and 0.85, same sign), giving f+ = 0.76 (95% CrI 0.42–0.94) and P(TM→JM) = 0.95. L658Q nearly eliminates the opposing strong JM→TM events, giving the cleanest bias of any condition: the activating mutation does not merely add TM→JM events but suppresses the counter-directed ones, reinstating the directed coupling of the signaling-competent state under ordered packing.

### FGFR3 A391E reproduces the pattern in a second receptor family

To test generality we applied the identical analysis to FGFR3 (PDB 2LZL), whose TM domain shares under 20% identity with EGFR and which carries the pathogenic activating A391E substitution (Crouzon syndrome with acanthosis nigricans); CASCADE resolved the same TM–JM coupling skeleton (Figure S4), to which PERI then assigned direction at 0.1 ns. The FGFR3 wild-type fluid dimer carried no net direction (two independent trajectories, f+ = 0.46 and 0.59, on opposite sides of 0.5; pooled f+ = 0.53, 95% CrI 0.39–0.67, P(TM→JM) = 0.65), balanced like the EGFR wild-type ordered dimer; unlike EGFR, which is constitutively active in fluid membranes, FGFR3 has no reported constitutive-activity phenotype in fluid, so this absence of a directed coupling is consistent with the directivity tracking the activity state of the receptor rather than the membrane phase alone. The ligand-dependent FGFR3 wild-type ordered dimer showed no resolved direction: CASCADE detects its TM–JM coupling from the many dimer frames (Figure S4), but PERI’s directed current is a relaxation transient that follows dimer formation (Figure 4b), and this dimer remains associated almost continuously, providing too few discrete association events to estimate f+. This reflects the physics rather than a limitation of the estimator: a continuously associated dimer is largely settled, so its fluctuations approach time-reversibility (detailed balance) and carry little net circulation, as for the balanced ligand-dependent EGFR ordered wild type. We therefore report it as n/a rather than force an estimate. The activating A391E mutant behaved like EGFR L658Q: both independent trajectories are TM→JM-biased (f+ = 0.74 and 0.78, same sign), giving f+ = 0.75 (95% CrI 0.54–0.89) and P(TM→JM) = 0.99. Thus in two receptor families whose TM domains share under 20% identity, an activating TM mutation produces a TM→JM directivity with high posterior probability (0.95 and 0.99).

### The coupling and its membrane precondition survive in all-atom simulation

To test force-field dependence, CASCADE was applied to an all-atom (CHARMM36m) simulation of the EGFR TM–JM dimer in the ordered membrane, backmapped from the coarse-grained trajectory at 10 μs and run for 5.8 μs without any secondary-structure restraint. The all-atom network reproduced the TM– JM coupling and a membrane-order coupling: local Scd was coupled to four of the seven protein variables (three JM and one TM; bootstrap frequency 0.52–0.91; Figure S3), fewer than in the coarse-grained ordered network, where Scd couples to all seven. The transmembrane helices remained folded throughout the all-atom trajectory (mean α-helical content 54 ± 2 residues by DSSP; Supporting Information). Because the all-atom dimer remains continuously associated (a single association episode), it does not yield the discrete association episodes the episode-based PERI analysis requires, so it validates the coupling detection only, and PERI direction is not applied to it. Because the all-atom model carries no elastic network or secondary-structure restraint of any kind, its reproduction of the coupling, together with the invariance of the PERI direction to the restraint scheme in the coarse-grained model (Methods; SI Note S4.1), indicates that the measured directivity reflects the physics of the membrane-embedded dimer rather than the coarse-grained model or its restraints.

### Summary across conditions

Across both receptor families the condition-level directivity at 0.1 ns tracks the activity state (Table 1, Figure 3): the activating TM mutants EGFR L658Q and FGFR3 A391E are TM→JM with posterior probability 0.95 and 0.99 (f+ = 0.76 and 0.75); wild-type EGFR in fluid membranes leans the same way (f+ = 0.60, P(TM→JM) = 0.93); and the ligand-dependent ordered wild type is balanced (f+ = 0.48, P(TM→JM) = 0.45). Every active-state trajectory shares the TM→JM sign, and the separation is robust to the threshold used to define a strong event (SI).

**Table 1.** Conformational directivity at 0.1 ns. Directivity is the net imbalance of the per-episode circulation between strong TM→JM and strong JM→TM association events; f+ = Σ(positive net)/Σ|net| is the TM→JM fraction of the directed mass (0.5 = balanced). f+, its 95% credible interval (CrI), and the posterior probability P(TM→JM) = P(f+ > 0.5) are obtained from a hierarchical Bayesian model over the independent trajectories (SI Note S3.4). The activating mutants (L658Q, A391E) are TM→JM with posterior probability 0.95 and 0.99; fluid wild type leans TM→JM (0.93); the ordered wild type is balanced (0.45). FGFR3 wild-type ordered is underpowered (near-continuous association, few episodes); FGFR3 wild-type fluid carries no net direction (balanced), consistent with the absence of a reported constitutive-activity phenotype for FGFR3 in fluid membranes.

| Condition | State | 0.1 ns trajectories (episodes) | $f_+$ (95% CrI) | $P(\text{TM} \rightarrow \text{JM})$ | Direction |
| --- | --- | --- | --- | --- | --- |
| EGFR wild type, fluid | active | 2 (60, 79) | 0.60 (0.46–0.73) | 0.93 | TM→JM |
| EGFR wild type, ordered | ligand-dependent | 2 (11, 10) | 0.48 (0.22–0.77) | 0.45 | none (balanced) |
| EGFR L658Q, ordered | active | 2 (6, 13) | 0.76 (0.42–0.94) | 0.95 | TM→JM |
| FGFR3 wild type, fluid | not established | 2 (77, 85) | 0.53 (0.39–0.67) | 0.65 | none (balanced) |
| FGFR3 wild type, ordered | ligand-dependent | limited (see text) | n/a | n/a | none (underpowered) |
| FGFR3 A391E, ordered | active | 2 (16, 16) | 0.75 (0.54–0.89) | 0.99 | TM→JM |

We tested the contrasts directly with the hierarchical Bayesian model (a two-level Bayesian bootstrap over trajectories and episodes; SI Note S3.4, Figure 5), which expresses each result as a posterior probability. The activating TM mutants are TM→JM with posterior probability 0.95 (L658Q) and 0.99 (A391E), and the ordered wild type is balanced (P(TM→JM) = 0.45). For the central contrast, the posterior probability that an activating TM mutation increases the TM→JM bias over the ordered wild type is 0.94; the corresponding probability for the fluid–ordered difference is 0.77. This is reinforced by simple reproducibility at the level of the independent trajectory, the unit of replication: all six active-state trajectories (two each for fluid wild type, L658Q and A391E) share the TM→JM sign, whereas the two ordered trajectories are split (Figure S5), exactly the pattern expected when the active states carry a directed coupling and the ligand-dependent state does not. The remaining uncertainty is dominated by the sampling of the ordered state, which associates only intermittently and yields few episodes; additional ordered trajectories will tighten the contrast.

**Figure 5.**
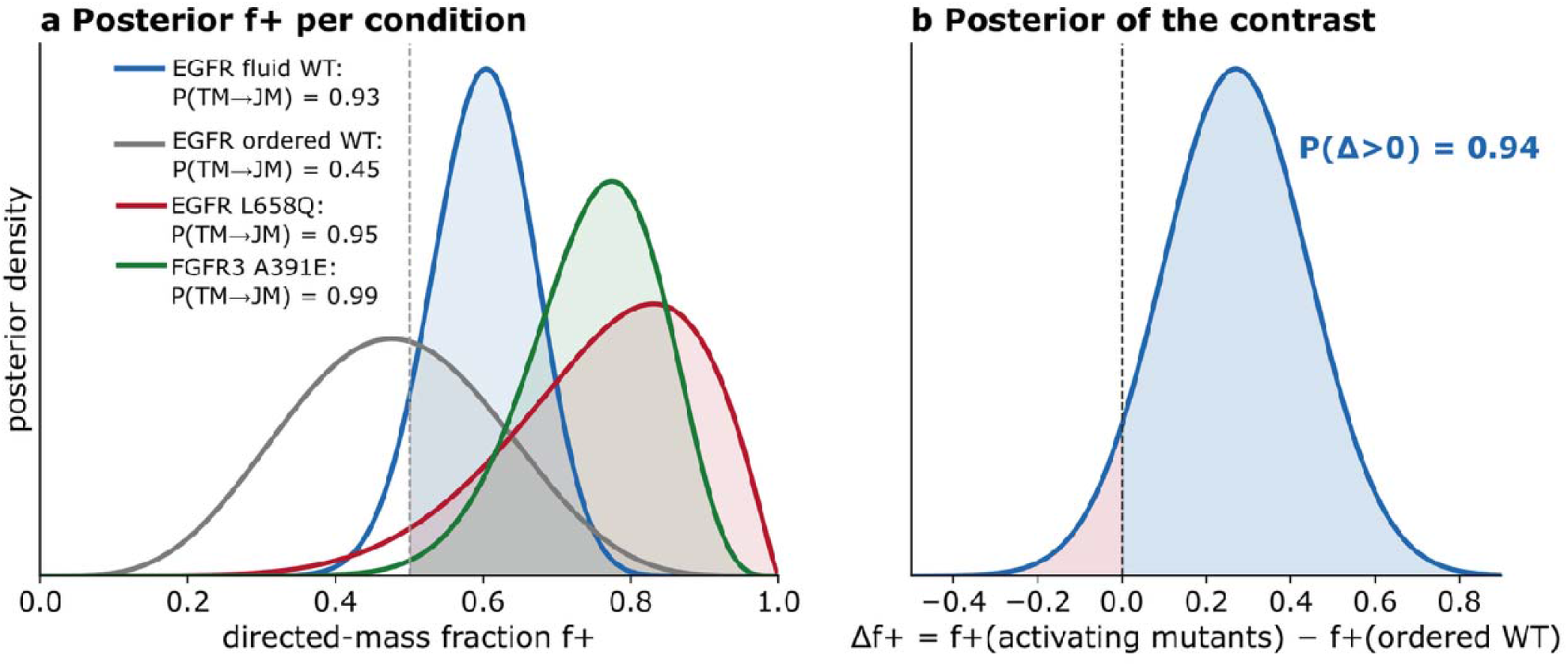
Hierarchical Bayesian directivity (two-level Bayesian bootstrap, 0.1 ns). (a) Posterior distribution of the directed-mass fraction f+ for each condition; the fraction of mass above 0.5 is P(TM→JM). The activating mutants and fluid wild type lie above 0.5; the ordered wild type straddles it. (b) Posterior of the contrast Δf+ = f+(activating mutants) − f+(ordered wild type): 94% of the posterior mass is positive, so P(an activating mutation increases the TM→JM bias) = 0.94.

## Discussion

The central contribution is that structural fluctuations within RTK dimers carry a measurable direction, that this direction distinguishes constitutively active from ligand-dependent states, and that the membrane environment and disease-associated mutations bias it (the activating-mutant directivities have posterior probability 0.95–0.99; the mutation increases the bias over the ordered wild type with posterior probability 0.94). Conformational directivity is a property of any multi-domain protein whose domains are coupled: it measures not which conformation a domain adopts but which of two coupled domains leads the other. It is inaccessible to structure alone and to methods that report coupling strength without direction.

At 0.1 ns the biological picture is a suppression of a bias, not an inversion of it. In every condition the dimer produces strong directed association events in both senses; what the membrane environment and the activating mutations set is their net balance. In the constitutively active states (wild-type EGFR in fluid membranes and the activating mutants) the strong TM→JM events outweigh the strong JM→TM events (condition-level directed-mass fraction f+ ≈ 0.60–0.76; posterior P(TM→JM) 0.93–0.99), giving a net fast TM→JM coupling; in the states without a net direction (the ligand-dependent ordered wild type, and wild-type FGFR3 in fluid, which has no reported constitutive-activity phenotype) the two senses equalize (f+ ≈ 0.5) and the net direction vanishes. A nonzero net directed coupling is, by construction, the signature of a dimer that has not settled, one still relaxing toward its stable configuration with a directional bias (broken detailed balance); the ordered environment does not invert this fast bias but removes it. Whether a net fast-timescale direction is present therefore reports whether the intramolecular coupling is settled or still descending its landscape with a preferred sense (Figure 6), consistent with the measured activity states of reconstituted EGFR.

**Figure 6.**
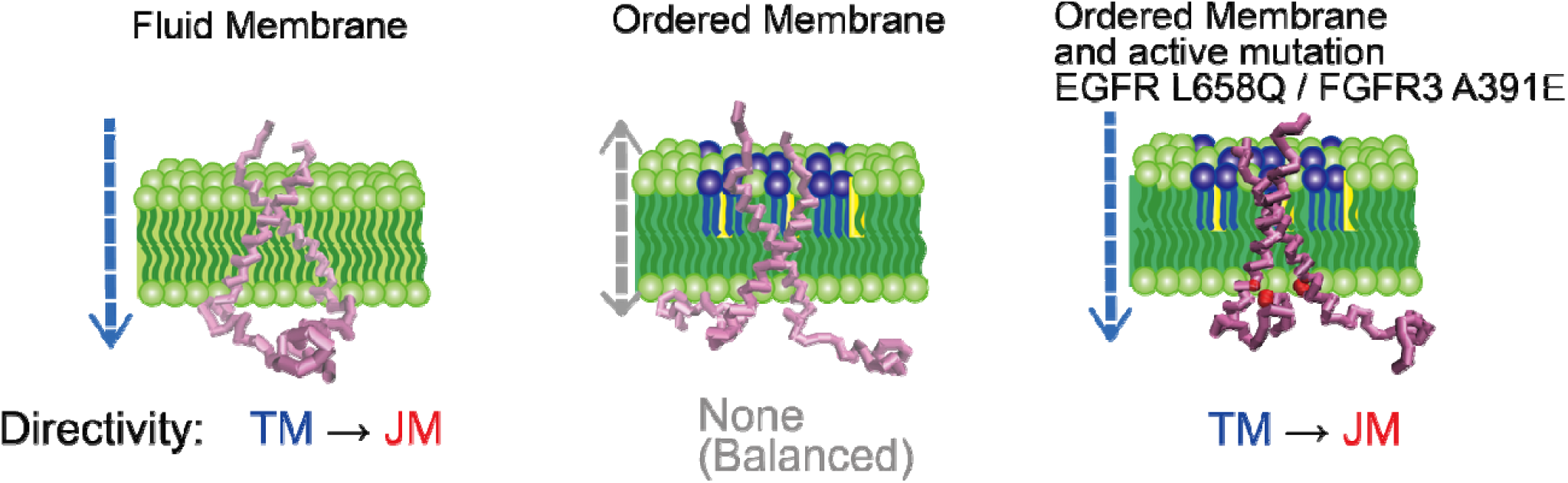
Activating mutations and the membrane environment set the directed TM→JM coupling. In wild-type EGFR in fluid membranes and under an activating TM mutation (EGFR L658Q, FGFR3 A391E), the dimer carries a net TM→JM coupling; in the ligand-dependent states (the ordered wild-type membrane, and wild-type FGFR3 in fluid) the two senses balance and no net direction remains. Th ordered environment does not reverse the coupling but equalizes it; activating TM mutations restore the TM→JM bias.

Two directed currents coexist at different timescales. A frequency decomposition of the phase-plane circulation separates a fast, sub-nanosecond TM→JM rotation from a slower, multi-nanosecond JM→TM rotation (Figure 4a). The fast rotation is the direct intramolecular coupling between the two covalently connected ends of one peptide; the slow rotation is a distinct, slower process, plausibly a membrane-mediated pathway in which the bilayer association of the JM drives the TM tilt on the lipid-reorganization timescale, although its molecular origin remains to be established. Sampling at 1 ns lies below the Nyquist limit of the fast rotation, so the circulation computed on 1 ns samples reads the slow JM→TM component, whereas 0.1 ns resolves the fast intramolecular TM→JM coupling, the one that defines the TM–JM relationship within a dimer, which dominates the full-bandwidth net. The two rotations are real and opposite, separated in time rather than in kind.

It is not obvious how a sub-nanosecond property can mark a signaling state established on the microsecond-to-millisecond timescale of kinase activation. We interpret the fast TM→JM directivity not as the activation event but as its mechanical precondition: a dimer in which TM fluctuations are efficiently and directionally transmitted to the JM is mechanically poised for the slower activating rearrangement, whereas one in which the fast transmission is balanced is not. The sub-nanosecond directivity is thus a readout of the mechanical predisposition of the dimer, a fast property that conditions a slow transition; its correspondence with the reconstitution activity states is, at this stage, an empirical one that full-length, longer simulations will test directly.

The directional assignment is consistent with, and gives a dynamical basis for, the long-standing structural model in which TM-helix rotation is propagated to the JM segment on activation.^6, 10^ That the activating TM substitution EGFR L658Q^22^ restores the TM→JM bias under ordered packing, and does so largely by suppressing the counter-directed events, indicates that the TM interface itself sets the directed coupling, and that pathogenic TM mutations act, at least in part, by reinstating it. The homologous FGFR3 A391E mutation (Crouzon syndrome with acanthosis nigricans)^23^ reproduces this at 0.1 ns in both independent trajectories (P(TM→JM) = 0.99); FGFR3, whose TM domain shares under 20% identity with EGFR, thus confirms that the association of a directed TM→JM coupling with an activating TM mutation is not idiosyncratic to one sequence.

One methodological point concerns how these results should be read in a membrane-protein context. We summarize direction by posterior probabilities rather than by a single significance threshold, because independent microsecond trajectories are costly and the ligand-dependent ordered dimer associates only intermittently, so its f+ posterior is wide; the limiting factor is the sampling of that state, not the size of the effect. The conclusion rests on a convergence: the activating mutants are TM→JM with posterior probability 0.95–0.99; the posterior probability that an activating mutation increases the bias over the ordered wild type is 0.94; the same TM→JM sign appears in all six active-state trajectories across two receptor families sharing under 20% TM identity; the mutations act specifically by suppressing the counter-directed events; and the pattern matches the independently measured ligand-dependence of these receptors.^5^ This follows current statistical practice, not to dichotomize on a single threshold but to weigh the totality and reproducibility of the evidence.^28^ Additional ordered-membrane trajectories will further tighten the mutation/membrane contrast.

The measured direction is not an artifact of the simulation restraints. No elastic network is applied: the only restraints on the protein are the standard MARTINI 2.2 secondary-structure-dependent backbone bonded terms, which are strictly intrachain and local, are interrupted at the TM–JM hinge, and act on no inter-chain geometry, so no bonded term acts on the very degrees of freedom (crossing angle, inter-TM distance, JM–JM distance and the lateral JM displacements) that carry the directional signal. The restraint scheme is moreover identical across the fluid and ordered membranes and across wild type and mutant, so it cannot generate a difference between conditions. Consistently, the all-atom CHARMM36m simulation, which carries no secondary-structure restraint of any kind, reproduces the same CASCADE coupling (PERI direction is not applied to the all-atom trajectory, because the dimer remains continuously associated (a single episode) and the episode-based analysis requires discrete association events; it validates the coupling detection only). Temporal resolution is equally essential, and reflects the scale-dependence of the coupling: the coupling has a fast, sub-nanosecond TM→JM rotation and a slower JM→TM one (SI Note S3), so at 1 ns the fast component is aliased and the slow JM→TM rotation dominates the apparent sign. We report the fast component because it is the direct mechanical coupling between the two covalently connected ends of one peptide, the intramolecular TM–JM coupling under study, whereas the slow rotation is a separate, slower relaxation; 0.1 ns sampling resolves it. Which component a given sampling returns is set by the timescale, not by statistical power.

Three limitations merit emphasis. First, we analyze TM–JM peptide dimers rather than intact receptors; whether the extracellular and kinase domains modulate the directivity will require full-length simulations, and the mechanistic link between a bound conformation and the directed coupling remains to be worked out.^6, 29^

Second, we do not measure signaling output directly; the association of TM→JM directivity with signaling competence is inferred from correspondence with reconstitution experiments,^5^ not measured. A third limitation is intrinsic to the measure: it applies only to dimers that undergo association dynamics, because the directed current is a relaxation transient that follows dimer formation. A continuously associated, settled dimer approaches detailed balance and carries little net current, so no direction can be assigned to it (as for wild-type FGFR3 in the ordered membrane), and the activity of such a persistently bound state cannot be predicted by this analysis. Within these limits, directivity analysis offers a route to predict the functional consequence of uncharacterized TM variants (an activating variant is predicted to restore TM→JM directivity) and, more broadly, a way to read a directed, membrane-controlled property of multi-domain membrane proteins that structure alone does not reveal.

## Methods

### Molecular dynamics

Coarse-grained simulations used GROMACS 2023.3^25^ with the MARTINI 2.2 force field^26, 27^, MARTINI polarizable water, 0.15 M NaCl, 20 fs timestep, velocity-rescaling thermostat (298 K), Parrinello– Rahman barostat (1 bar), reaction-field electrostatics (relative permittivity 2.5, Coulomb cutoff 1.1 nm), and a van der Waals cutoff of 1.1 nm with a potential-shift modifier. Inputs were generated with CHARMM-GUI Martini Maker^30^ and its standard production parameters used unchanged. Protein structures were converted to the MARTINI representation with martinize (v2.5) using the martini22p parameter set, with secondary structure assigned by DSSP^31^. No elastic network was applied: secondary structure is maintained by the standard MARTINI 2.2 secondary-structure-dependent backbone bonded terms alone (helical backbone bonds and angles, and i→i+3 backbone dihedrals, φ□ = −120°, k = 400 kJ mol□^1^). The topology therefore contains no distance restraint derived from a reference structure: the longest-range bonded term is the i→i+3 helical dihedral, the backbone dihedral restraints are interrupted at the TM–JM hinge, and no term acts on inter-chain geometry. Backbone position restraints exist only inside an #ifdef POSRES block (CHARMM-GUI, for equilibration) and are inactive in production. EGFR TM–JM: residues 634–683 (TM helix 646–668; intracellular JM 669–683) from PDB 2M20. FGFR3 TM–JM: corresponding segment from PDB 2LZL. Mutations built with CHARMM-GUI. Fluid membrane (8 × 8 × 15 nm^3^): DIPC (70 mol%)/DOPS (30 mol%). Ordered membrane (40 × 40 × 15 nm^3^): upper leaflet DIPC/DPSM/CHOL equimolar, lower leaflet DIPC/CHOL/DOPS equimolar; four peptide copies per system (the fluid systems used two), initial separation 5 nm. Dimer pairs and association episodes were defined in two steps: in multi-peptide boxes the dimer pair is first identified by the TM center-of-mass distance (a pair below 20 Å is a dimer; the single longest-lived pair enters each analysis set), and within that pair association episodes are then defined by the TM backbone minimum distance < 6 Å sustained ≥ 100 ns. Episodes shorter than 100 frames (10 ns at 0.1 ns sampling) are excluded from the circulation analysis. A single dimer pair therefore enters each analysis set, so an analysis set is not a mixture of two dimer configurations (SI Note S4.3); the EGFR ordered trajectories each contributed one pair (PROA–PROD, PROA–PROC). All-atom validation: a CG ordered-membrane snapshot at 10 μs was backmapped to CHARMM36m, which carries no secondary-structure restraint of any kind, and run 5.8 μs. Trajectories were saved every 100 ps; for the direction analysis, structural observables were extracted at 0.1 ns (see below).

### Coupling detection (CASCADE)

CASCADE identifies co-fluctuating variable pairs: hill-climbing DAG search with Bayesian information scoring^32^ (10 restarts), bootstrap edge stability (n = 500, consensus ≥ 0.50) with false-discovery-rate control^33^, and Bayesian structural-equation effect sizes (PyMC^20^, No-U-Turn sampler^34^, standardized variables). A variable with consensus edges to ≥ 80% of others is treated as a dominant environmental regulator (local membrane order, Scd); its effect is regressed out and the pipeline rerun on residuals to expose internal protein coupling. CASCADE is used here only to detect coupling (the undirected skeleton); direction is assigned by the circulation analysis.

### Direction (PERI: phase-plane circulation)

PERI (Phase-plane Estimation of Rotational Irreversibility) assigns direction as follows. For each pre-registered TM–JM variable pair (x = TM, y = JM), standardized within each association episode, the per-step signed area c(t) = x_mΔy − y_mΔx (midpoints x_m, y_m; one-step differences) is summed over the episode and over the pairs; positive = TM→JM. This equals the net probability current in the (x, y) plane and is nonzero only when detailed balance is broken. Significance is assessed against a within-episode circular-shift null (JM series rolled by a random offset per episode, wrap step dropped) that preserves the distribution, autocorrelation and nonlinearity of each series while destroying TM–JM temporal alignment. The estimator was validated on synthetic AR residuals with an injected lead (recovers TM→JM at z ≈ +9.8; null on simultaneous coupling; false-positive rate at nominal, including under differing smoothness and a common driver).

### Temporal resolution

The directed coupling is scale-dependent (SI Note S3): a fast, sub-nanosecond TM→JM rotation and a slower JM→TM one, so on identical episodes the apparent net inverts between 1 ns (which reads the slow component) and 0.1 ns (which resolves the fast component). Structural observables were therefore extracted at 0.1 ns to resolve the fast intramolecular TM–JM component. The membrane-order regression is not required for direction (the raw 0.1 ns geometry and the membrane-regressed residuals give identical circulation), so 0.1 ns extraction computes only the geometric observables.

### Statistical replication

The independent trajectory is the unit of replication. Direction is estimated per independent trajectory and pooled across independent trajectories of the same condition; the number of independent trajectories, the per-trajectory direction, and their sign agreement are reported. Within-trajectory significance addresses whether a given simulation shows a direction; the generalizable claim for a condition requires agreement across independent trajectories.

### Table of symbols and acronyms

**Table 2.** Symbols and acronyms used in the main text and figures.

| Symbol / term | Meaning |
| --- | --- |
| TM | transmembrane domain |
| JM | intracellular juxtamembrane segment |
| Scd | local lipid acyl-chain order parameter (membrane order around a chain) |
| tilt_a, tilt_b | tilt angle of TM helix A / B relative to the membrane normal |
| crossing_angle | angle between the two TM helices |
| inter_tm_distance | distance between the TM helix centers |
| jm_a_x, jm_b_x | lateral (membrane-plane) displacement of the intracellular JM of chain A / B |
| jm_jm_distance | distance between the two intracellular JM segments |
| circulation | net signed area swept per unit time in the (TM, JM) plane = net probability current (+ = TM leads JM) |
| directivity | the net direction of causal influence between coupled structural fluctuations |
| CASCADE | Bayesian-network structural-covariance framework, used here for coupling detection (edge orientation not interpreted as direction; direction is assigned by PERI) |

## Data and Code Availability

Trajectories are available upon request. CASCADE: github.com/takeshi-sato-dev/cascade. PERI (phase-plane circulation) directivity code and synthetic validation: github.com/takeshi-sato-dev/peri (deposited on acceptance).

AI Disclosure. Claude (Anthropic) was used for programming assistance and language editing. All methodological design, validation strategy, and scientific interpretation are the authors’ own.

## ASSOCIATED CONTENT

### Supporting Information

The Supporting Information is available free of charge at https://pubs.acs.org/doi/XXXX.

Full mathematical framework for CASCADE and PERI; synthetic-data validation; hierarchical Bayesian model; stationarity and sensitivity analyses; all-atom validation and transmembrane-helix folding analysis; molecular dynamics and analysis details; Supporting Figures and Tables (PDF).

## AUTHOR INFORMATION

### Author

Hiroko Tamagaki-Asahina — Kyoto Pharmaceutical University, Kyoto 607-8414, Japan; ORCID:

## Notes

The authors declare no competing financial interest.

## ACKNOWLEDGMENT

This work was supported by the Kyoto Pharmaceutical University Fund for the Promotion of Collaborative Research.

## TOC

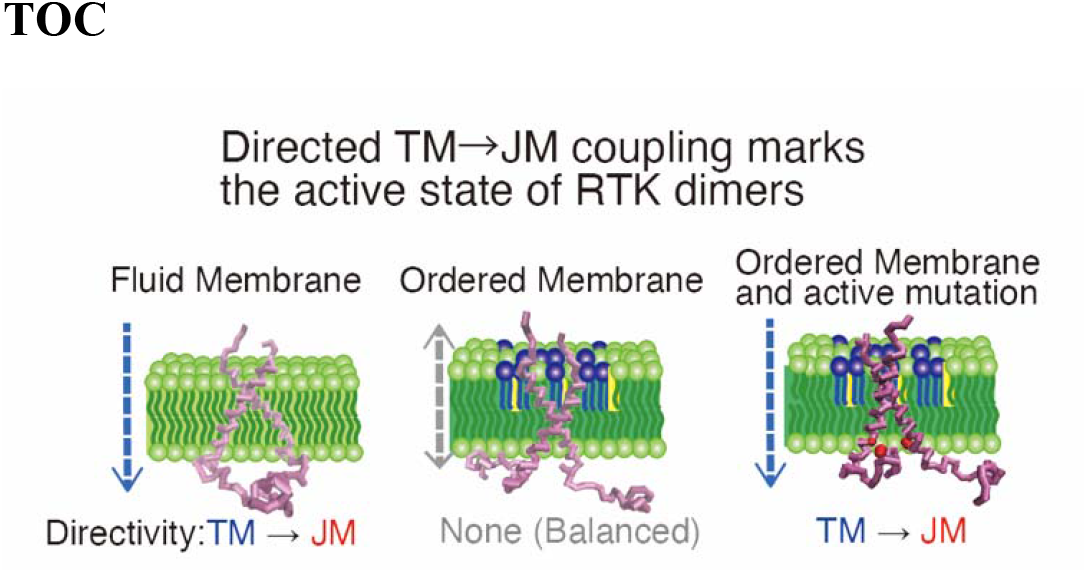

## Notes

https://github.com/takeshi-sato-dev/cascade

https://doi.org/10.5281/zenodo.18626137

https://doi.org/10.5281/zenodo.18626075

https://doi.org/10.5281/zenodo.19127206

https://doi.org/10.5281/zenodo.19412949

### Competing Interest Statement

The authors have declared no competing interest.

### Summary of Updates

This is a revised version of the manuscript. The title has been changed. The method presented in version 1 under the name MERIT is presented here under the name PERI (Phase-plane Estimation of Rotational Irreversibility). The figures and the Supporting Information have been revised accordingly.

