## Supporting Information for "A directed TM→JM coupling in receptor tyrosine kinase dimers, set by activating mutations and the membrane environment"

*This Supplementary Information specifies, in full, the two analytical components of the study: CASCADE, which detects inter-domain coupling, and the phase-plane circulation estimator, which measures the direction of that coupling. It also documents the scale-dependence of the directed coupling (a fast TM→JM and a slow JM→TM rotation) that mandates 0.1 ns sampling, the Bayesian treatment of inference and replication, and robustness controls.*

### **Abbreviations**

Abbreviations used in the manuscript and this Supporting Information (the structural-variable glossary is main-text Table 2).

| **Abbreviation / symbol** | **Definition** |
| --- | --- |
| RTK | receptor tyrosine kinase |
| TM | transmembrane domain |
| JM | intracellular juxtamembrane segment |
| MD | molecular dynamics |
| CG | coarse-grained |
| AA | all-atom |
| NMR | nuclear magnetic resonance |
| ATR-IR | attenuated total reflectance infrared spectroscopy |
| PCA | principal component analysis |
| MI | mutual information |
| DAG | directed acyclic graph |
| BIC | Bayesian information criterion |
| SEM | structural-equation model |
| NUTS | No-U-Turn sampler |
| FDR | false-discovery rate |
| FPR | false-positive rate |
| CrI | credible interval (Bayesian posterior interval) |
| CI | confidence interval (frequentist) |
| CASCADE | the Bayesian-network structural-covariance framework used here for coupling detection |
| PERI | Phase-plane Estimation of Rotational Irreversibility (the direction measure) |
| Scd | local lipid acyl-chain order parameter (membrane order) |
| f+ | directed-mass fraction: the TM→JM share of the directed circulation (0.5 = balanced) |
| DIPC | dilinoleoyl phosphatidylcholine |
| DOPS | dioleoyl phosphatidylserine |
| DPSM | palmitoyl sphingomyelin |
| CHOL | cholesterol |
| GM3 | monosialodihexosylganglioside GM3 |

### **Supplementary Note S1. CASCADE: detection of inter-domain coupling**

CASCADE, a Bayesian-network structural-covariance framework, identifies which structural variables co-fluctuate. In this study it is used to (i) establish the undirected coupling skeleton (which TM and JM variables are statistically dependent) and (ii) remove the dominant membrane-order signal so that internal protein coupling is exposed. Edge orientation is not interpreted as direction; direction is assigned separately by PERI (Note S2).

#### **S1.1 Bayesian-network model**

The joint distribution of the p structural variables is modeled as a Bayesian network, a directed acyclic graph (DAG) in which each node is a variable and each edge encodes a conditional dependence. The distribution factorizes over the graph as

*P(X₁, …, Xₚ) = ∏ᵢ P(Xᵢ | Pa(Xᵢ)),*

where Pa(Xᵢ) is the parent set of Xᵢ. Structure learning seeks the DAG that best explains the observed frame × variable matrix under a penalized-likelihood score.

#### **S1.2 Scoring function**

Candidate DAGs are scored by the Gaussian Bayesian Information Criterion. For a node Xᵢ with parent set Pa(Xᵢ),

*BIC(Xᵢ | Pa(Xᵢ)) = n · ln(RSSᵢ / n) + kᵢ · ln(n),*

with n the number of frames, RSSᵢ the residual sum of squares from the linear regression of Xᵢ on Pa(Xᵢ), and kᵢ = |Pa(Xᵢ)| + 1 the parameter count (coefficients plus intercept). The total score is the sum of local scores over all nodes; lower is better. The first term rewards fit, the second penalizes complexity; the penalty is essential for MD data, where the structural variables of covalently connected regions are pervasively correlated and an unpenalized score would return a fully connected, uninformative graph.

#### **S1.3 Structure search**

The score is maximized by hill-climbing over DAG space. From the empty graph, at each step the three local operations (edge addition, deletion, and reversal) are evaluated and the one giving the largest score improvement is applied, subject to an acyclicity constraint (any operation creating a directed cycle is rejected); the search terminates when no operation improves the score, and is repeated from 10 random restarts. A constraint-based (PC) alternative was rejected because it oriented all 42 tested edges at frequency 1.0 on the EGFR fluid data, providing no discrimination between strong and weak relationships.

#### **S1.4 Bootstrap edge stability**

Edge reliability is assessed by nonparametric bootstrap: frames are resampled with replacement (500 resamples for the protein trajectories, 100 for the synthetic benchmarks) and the full hill-climbing search is repeated on each. The bootstrap frequency of an edge is the fraction of resamples in which it appears in the optimal DAG; the consensus network retains edges with frequency ≥ 0.50. Varying the threshold over {0.40, 0.50, 0.60} leaves the hierarchy direction, root-node identity, and effect-sign pattern unchanged.

#### **S1.5 Effect quantification (Bayesian structural-equation model)**

For each node Yᵢ with consensus parents Xⱼ, standardized effect sizes are estimated from the structural-equation model

*Yᵢ = α + Σⱼ βⱼ Xⱼ + ε, ε ~ Normal(0, σ²),*

with all variables standardized (zero mean, unit variance) so that the βⱼ are comparable across units (Å, degrees, dimensionless Scd). Posteriors are sampled with PyMC (Normal(0,1) priors on βⱼ, HalfNormal(1) on σ; NUTS, 4 chains × 2,000 draws, 1,000 tuning; R̂ < 1.01), giving each edge a posterior mean effect and 95% credible interval.

#### **S1.6 Hierarchical removal of the membrane-order signal**

A variable D is classified as a dominant environmental regulator if it has consensus edges (bootstrap frequency > 0.50) to at least 80% of the other variables, a hub signature statistically distinct from pairwise correlation. In the ordered systems the local membrane-order parameters Scd meet this criterion. Its variance is then removed from every remaining variable before the internal coupling is read out:

(1) for each remaining variable Yⱼ, regress on the dominant set D₁, …, Dₘ by ordinary least squares; (2) form the residual

*rⱼ = Yⱼ − Ŷⱼ(D₁, …, Dₘ);*

(3) rerun the full CASCADE pipeline (hill-climbing, bootstrap, Bayesian SEM) on the residual matrix {rⱼ}. The decomposition is applied identically to every condition, so all conditions enter the direction analysis with the same preprocessing. The membrane regression is required only for the coupling read-out; the PERI direction estimator is insensitive to it (the circulation on the raw geometry equals that on the residuals; Note S3.3).

#### **S1.7 Identifiability limit (why CASCADE is not used for direction)**

Because BIC depends on the covariance and not on time order, the orientation of an edge is only partially identifiable: within a Markov-equivalence class, DAGs encoding the same conditional-independence relations receive identical scores (e.g. X→Y→Z and Z→Y→X are indistinguishable). On synthetic benchmarks CASCADE recovered every true edge (true-positive rate 1.00, false-positive rate 0.00 across chain, hub, and mixed topologies and sample sizes 500–4,000 frames), but orientation accuracy fell to 50–75% for chain motifs, precisely the TM–JM motif here. CASCADE is therefore used strictly as a coupling detector; direction is obtained from PERI (Note S2).

#### **S1.8 Structural variable definitions**

Nine observables per frame. Membrane order: local_scd_a / local_scd_b, the acyl-chain order parameter of lipids within 12 Å of each TM helix,

*S_cd = ½ ⟨ 3 cos²θ − 1 ⟩,*

θ the angle between consecutive tail-bead bond vectors and the bilayer normal, averaged over the local lipids. TM geometry: tilt_a / tilt_b, the tilt of each TM helix principal axis to the bilayer normal; crossing_angle, the angle between the two TM principal axes projected in the membrane plane; inter_tm_distance, the TM backbone center-of-mass separation. JM position (intracellular): jm_a_x / jm_b_x, the in-plane lateral displacement of each JM center of mass from the dimer center; jm_jm_distance, the distance between the two intracellular JM centers. A JM–JM contact count (inter-chain JM beads within 6 Å) is also computed; inter-JM contacts are sparse in these dimers (of order 1% of frames), so the continuous jm_jm_distance is used as the primary JM–JM variable.

### **Supplementary Note S2. PERI: phase-plane circulation for direction**

PERI (Phase-plane Estimation of Rotational Irreversibility) measures the direction of a detected TM–JM coupling as the net circulation of the joint state in the phase plane, and the idea is best seen as a picture. Represent the dimer at each instant by a single point whose horizontal coordinate is the TM state x and whose vertical coordinate is the JM state y; as the dimer fluctuates, this point moves and traces a path. If the two domains moved in perfect lockstep the point would slide back and forth along one diagonal line; it would retrace itself, enclose no area, and single out no leader. But if one domain systematically leads the other (say TM shifts first and JM follows a moment later), the point cannot retrace its path: it travels out along one route and returns along another, closing a loop. A loop has a sense of rotation, like the sweep of a clock hand, and it encloses an area. That sense of rotation is the direction of the coupling (one way of turning is TM→JM, the other JM→TM), and the enclosed area measures how strongly one domain leads. This persistent looping of the joint state, a rotational bias in how the two coordinates co-move rather than any flow of matter, is what we call the circulation.

Why a current, and why it must vanish at rest: at thermodynamic equilibrium every clockwise excursion is on average matched by a counter-clockwise one, the areas cancel, and there is no net loop. A persistent net loop can arise only while the system is being driven or is still relaxing, that is, only when its motion is not identical run forwards and backwards in time. Reversing the movie of the trajectory turns every loop the other way, so the net loop is exactly the part of the motion that distinguishes forward from backward time. This is why the net circulation is the net probability current and a direct measure of broken detailed balance (formalized in Note S2.2).

How to read the result: each association episode contributes one signed loop-area, positive for a TM→JM turn, negative for a JM→TM turn. Because the dimer produces strong loops of both senses, PERI does not average them but asks what fraction of the total loop area turns the TM→JM way: the directed-mass fraction f₊ (Note S2.3). Read it on a simple scale: f₊ = 0.5 is balanced (no leader), f₊ > 0.5 is a net TM→JM lead, f₊ < 0.5 a net JM→TM lead. A handful of strong, well-formed loops dominate f₊ while the many faint, nearly loop-free episodes count for almost nothing, so f₊ reports the direction of the strong association events, not of background jitter. Figure 3a shows one episode directly: an open, oriented loop for a directed episode versus a collapsed diagonal for an in-phase one.

#### **S2.1 The circulation statistic**

For a standardized TM–JM pair (x, y) sampled within an association episode of length T, the signed area swept per step is computed by the midpoint (trapezoidal) rule,

*c(t) = xₘ Δy − yₘ Δx, xₘ = ½(x_t + x_{t+1}), Δx = x_{t+1} − x_t,*

and the episode circulation is its sum,

*C = Σ_{t=1}^{T−1} c(t) = ∮ (x dy − y dx) = 2A,*

which, by Green’s theorem, equals twice the signed area A enclosed by the orbit in the (x, y) plane; equivalently C = Σ r² Δφ accumulates the rotational bias, with φ the phase angle. The condition statistic sums C over the pre-registered TM–JM pairs (the full 4 TM × 3 JM set) and over episodes. By convention C > 0 ⇔ TM→JM, C < 0 ⇔ JM→TM, C ≈ 0 ⇔ no directed coupling.

#### **S2.2 Net probability current and broken detailed balance**

For the coarse dynamics viewed as a stationary Markov / overdamped-Langevin process on (x, y), the mean per-step circulation equals the area integral of the curl of the stationary probability current J(x, y):

*⟨C⟩ = ∫∫ (∂ₓ J_y − ∂_y J_x) dx dy.*

At thermodynamic equilibrium detailed balance holds, the stationary current vanishes everywhere (J ≡ 0), and ⟨C⟩ = 0. A nonzero ⟨C⟩ therefore certifies broken detailed balance, a non-equilibrium state in which the dimer is still relaxing toward its stable configuration with a directional bias. Under time reversal t → −t the orbit is traversed backwards and c(t) → −c(t), so C is a time-reversal-odd observable: it is exactly the component of the dynamics that distinguishes forward from backward time, i.e. the physically correct signature of a directed process. A symmetric, time-reversible potential (including a harmonic elastic network) can shape the covariance but cannot produce a nonzero ⟨C⟩.

#### **S2.3 Directed-mass fraction**

Because the dimer emits strong directed events of both signs, a condition is summarized not by any single episode but by the imbalance of its episode circulations {C_e}. The threshold-free directed-mass fraction is

*f₊ = Σ_e max(C_e, 0) / Σ_e |C_e| ∈ [0, 1], f₊ = 0.5 ⇔ balanced (no net direction).*

Low-amplitude episodes contribute ≈ 0 to both sums and are down-weighted automatically, so f₊ is dominated by the strong directed events and is insensitive to the many near-zero episodes; f₊ > 0.5 is a TM→JM bias, f₊ < 0.5 a JM→TM bias.

#### **S2.4 Null model (estimator calibration) and replication**

The estimator is calibrated against a within-episode circular-shift null: within each episode the JM series y is rolled by a random offset, y_t → y_{(t+τ) mod T} (the single wrap step excluded), which preserves the marginal distribution, autocorrelation, and nonlinearity of y exactly while destroying only its temporal alignment with x. A nonzero net that survives this null is what a directed coupling (not spurious alignment or shared smoothness) produces; the standardized departure z = (net − mean_null)/sd_null is used as a calibration diagnostic. A block shift (one offset per episode applied to all JM variables) preserves the mutual JM–JM correlations. This null is what establishes that the estimator has a controlled false-positive rate (Note S2.5); the directional inference itself is Bayesian (Note S3.4).

The unit of replication is the independent trajectory, not the episode (episodes within one trajectory share lipid realization and starting configuration). Because a single trajectory yields few episodes, the direction of a condition is inferred at the condition level by the hierarchical Bayesian model of Note S3.4, which pools the independent trajectories and returns the posterior probability P(f₊ > 0.5) = P(TM→JM) and a 95% credible interval for f₊. Per-trajectory f₊ values are reported to show sign reproducibility across independent simulations.

#### **S2.5 Synthetic validation**

Injected circulations of known sign are recovered, and a simultaneous (in-phase) coupling returns no net, tested against a within-episode circular-shift null (reproducible, seed 0, via synthetic_validation/dir_current.py --controls). The same estimator computed on the linear lead–lag benchmark (dir_current_LINLAG.csv) returns a TM→JM current at z = +8.8, while CCM on that identical linear signal returns no direction (system_verdict_LINLAG.csv, z = 0.37); on a nonlinear (logistic) coupling CCM instead detects the direction (z = +5.5; SYN_logistic, ccm_disjoint_LOG.log), so CCM's blindness is specific to the linear lead–lag:

*Table S1 | Synthetic validation of PERI. On synthetic episodes with a known injected circulation, the pooled net (over the 4×3 TM–JM pairs) and its z-score against a within-episode circular-shift null recover the planted direction (per-pair z +8.4 to +9.7 for TM→JM, −8.9 to −9.7 for JM→TM); a simultaneous (in-phase) coupling returns no net. Values are the seed-0 output of synthetic_validation/dir_current.py --controls.*

| **Synthetic case** | **net (all pairs)** | **z (vs shift-null)** | **p** | **verdict** |
| --- | --- | --- | --- | --- |
| Injected TM→JM lead | +4248 | +9.8 | 0.003 | TM→JM (recovered) |
| Injected JM→TM lead | −4331 | −9.1 | 0.003 | JM→TM (recovered) |
| Simultaneous (in phase) | −167 | −0.3 | 0.36 | no direction (null) |

The false-positive rate under the null (independent series, no coupling) is controlled, including the adversarial cases of unequal smoothness and a shared slow driver:

*Table S2 | Calibrated false-positive rate of the circulation null (independent series, no coupling), by null configuration.*

| **Null configuration** | **FPR at α = 0.05** |
| --- | --- |
| independent, equal autocorrelation | 0.075 |
| independent, different autocorrelation (smoothness) | 0.013 |
| common slow driver, simultaneous, different noise | 0.000 |

The independent-equal-autocorrelation FPR is 0.075, marginally above the 0.05 nominal level; we report it explicitly and note that the condition-level bootstrap CI (Note S2.4), not a single-test threshold, governs every directional claim. The amplitude-normalized variant (Δφ/Δt) was rejected because it diverges near the origin and inflates the false-positive rate on simultaneous coupling.

#### **S2.6 Directivity as the net balance of bidirectional association events**

Within a single condition the per-episode circulation is not one-signed: the dimer produces strong directed events in both senses. We characterize a condition by the net imbalance of these events, summarized by the threshold-free directed-mass fraction f+ = Σ(positive per-episode net) / Σ|net|, the fraction of the total directed mass that is TM→JM (f+ = 0.5 = balanced, no net direction; f+ > 0.5 = TM→JM bias). Low-amplitude episodes contribute negligibly, so the measure is dominated by the strong events. Because a single trajectory yields few episodes, direction is inferred at the condition level (hierarchical Bayesian model, Note S3.4); the per-trajectory net and f+ are reported below to show sign reproducibility.

*Table S3 | Per-trajectory directivity at 0.1 ns: episodes, net circulation, and directed-mass fraction f+ for each independent trajectory.*

| **Trajectory (0.1 ns)** | **state** | **episodes** | **net** | **f+** |
| --- | --- | --- | --- | --- |
| EGFR fluid1 | active | 60 | +968 | 0.62 |
| EGFR fluid2 | active | 79 | +936 | 0.59 |
| EGFR L658Q rep2 | active | 13 | +345 | 0.85 |
| EGFR L658Q rep1 | active | 6 | +118 | 0.71 |
| FGFR3 A391E rep1 | active | 16 | +525 | 0.74 |
| FGFR3 A391E rep2 | active | 16 | +258 | 0.78 |
| EGFR ordered1 | ligand-dep | 11 | −43 | 0.42 |
| EGFR ordered2 | ligand-dep | 10 | +11 | 0.52 |
| FGFR3 fluid rep1 | not established | 77 | −187 | 0.46 |
| FGFR3 fluid rep2 | not established | 85 | +607 | 0.59 |

Pooled to the condition level (hierarchical Bayesian posterior, Note S3.4):

*Table S4 | Condition-level directivity at 0.1 ns, pooled over the independent trajectories (hierarchical Bayesian f+, 95% CrI, and P(TM→JM)).*

| **Condition** | **state** | **episodes** | **f+** | **95% CrI** | **P(TM→JM)** | **direction** |
| --- | --- | --- | --- | --- | --- | --- |
| EGFR fluid WT | active | 139 | 0.60 | 0.46–0.73 | 0.93 | TM→JM |
| EGFR L658Q | active | 19 | 0.76 | 0.42–0.94 | 0.95 | TM→JM |
| FGFR3 A391E | active | 32 | 0.75 | 0.54–0.89 | 0.99 | TM→JM |
| EGFR ordered WT | ligand-dep | 21 | 0.48 | 0.22–0.77 | 0.45 | balanced |
| FGFR3 fluid WT | not established | 162 | 0.53 | 0.39–0.67 | 0.65 | balanced |

All six active-state trajectories (two each for fluid wild type, L658Q, A391E) are TM→JM-biased (f+ = 0.59–0.85, all on the same side of 0.5); the two ordered wild-type trajectories are balanced (f+ = 0.42, 0.52) and disagree in the sign of their near-zero net, as expected for a genuine absence of direction; the two FGFR3 fluid wild-type trajectories are likewise balanced, falling on opposite sides of 0.5 (f+ = 0.46, 0.59) and pooling to f+ = 0.53 (P(TM→JM) = 0.65). At the condition level the activating mutants L658Q and A391E are TM→JM with posterior probability 0.95 and 0.99; fluid wild type leans TM→JM (0.93); ordered wild type is balanced (0.45). The separation is robust to the amplitude threshold used to define a strong event: over thresholds from 20 to 50 (per-episode net units) the ratio of strong TM→JM to strong JM→TM mass exceeds 1 for every active trajectory and is at or below 1 for the ordered trajectories.

**CASCADE coupling networks and edge statistics (all systems)**

CASCADE is used here only to detect coupling (the undirected skeleton; direction is measured by PERI). The following figures are the CASCADE coupling networks and their bootstrap edge statistics for the simulated systems, as reported in the original submission; the edge counts are unchanged (EGFR fluid 25 full / 10 internal; EGFR ordered 30 full / 15 internal; EGFR L658Q ordered 8 internal). The all-atom result validates CASCADE coupling only (the direction measure PERI is not applied to it). Over the 5.8 μs all-atom trajectory the two transmembrane helices remained folded, confirming the fold is maintained without any secondary-structure restraint: the DSSP α-helical content was 54 ± 2 residues (~27 per chain) and did not decline over the trajectory (52.8, 52.8 and 55.3 residues in successive thirds) (gmx dssp: Gorelov, S.; Titov, A.; Tolicheva, O.; Konevega, A.; Shvetsov, A. J. Chem. Inf. Model. 2024, 64, 3593–3598; Kabsch, W.; Sander, C. Biopolymers 1983, 22, 2577–2637).

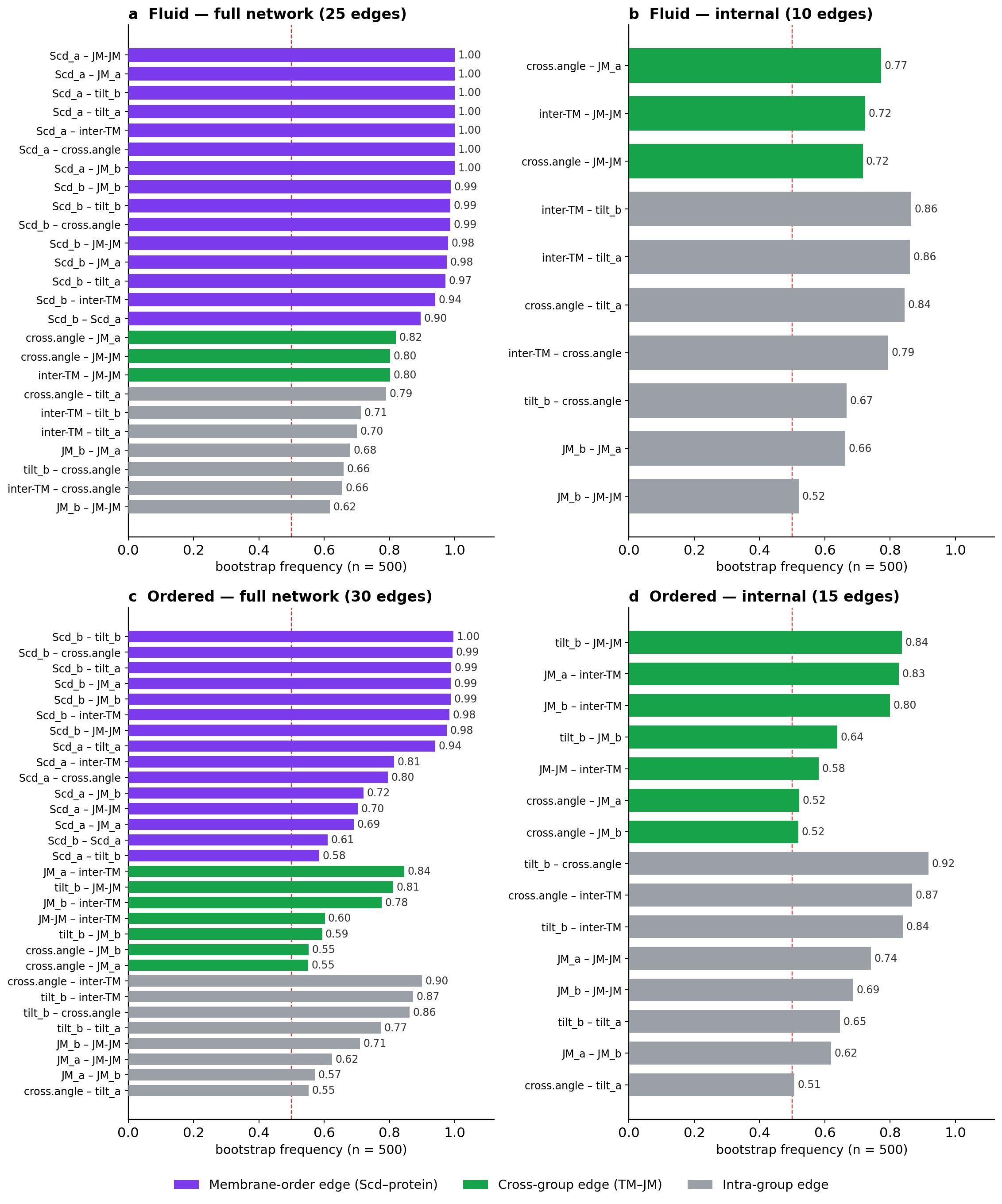

*Figure S1 | CASCADE bootstrap edge-frequency distributions (EGFR fluid and ordered membranes). Horizontal bar charts of edge bootstrap frequency (n = 500) with the 50% consensus threshold (dashed): (a) fluid full network (25 edges), (b) fluid internal network after Scd removal (10 edges), (c) ordered full network (30 edges), (d) ordered internal network after Scd removal (15 edges). Edges are shown undirected (orientation is not interpreted; Note S1.7); colors denote edge type (membrane-order Scd–protein, cross-group TM–JM, intra-group).*

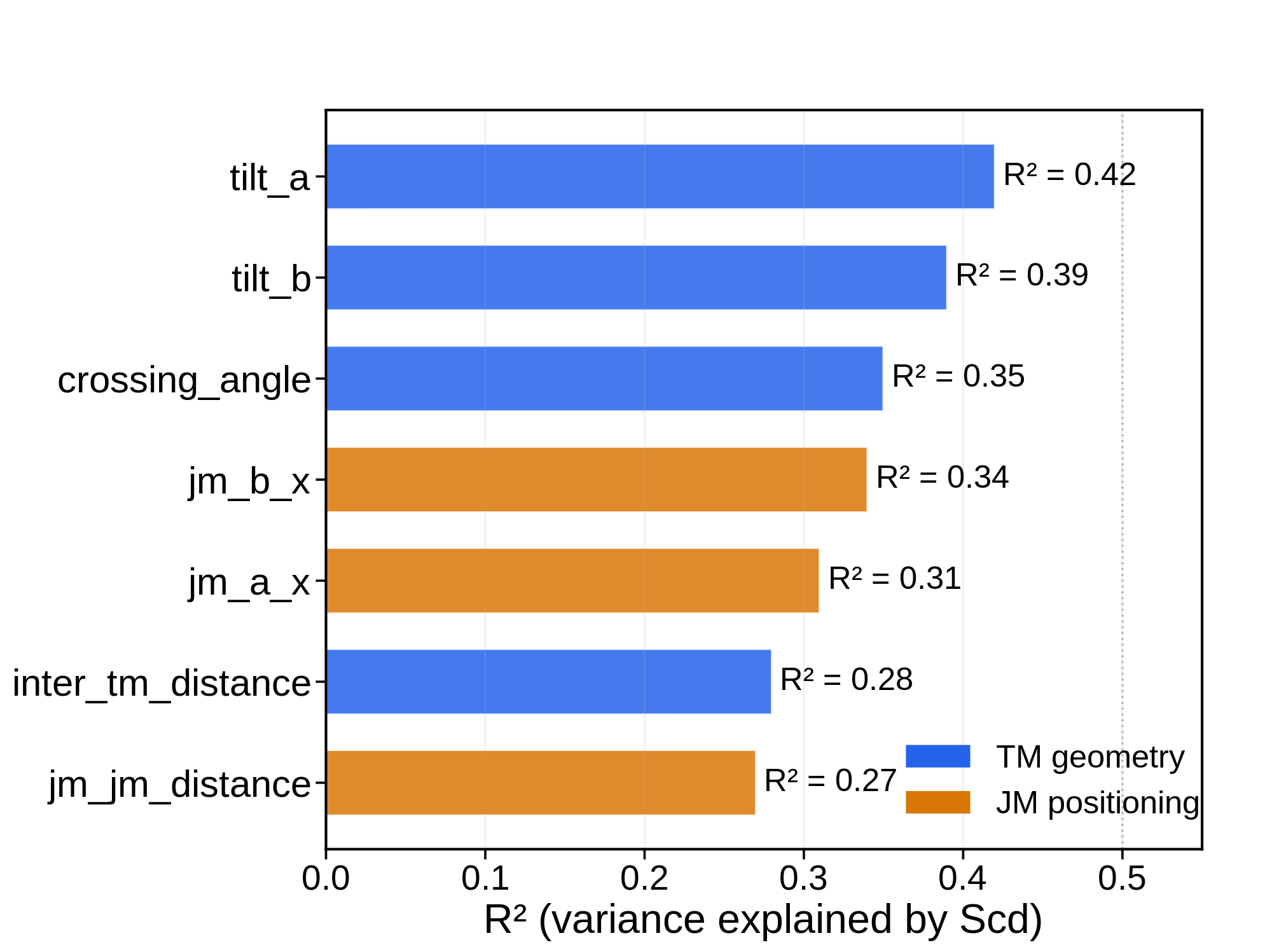

*Figure S2 | Hierarchical-analysis R² for the EGFR fluid membrane. Fraction of variance in each structural variable explained by the membrane-order (Scd) variables, motivating the hierarchical removal of the Scd variance before the internal protein network is read.*

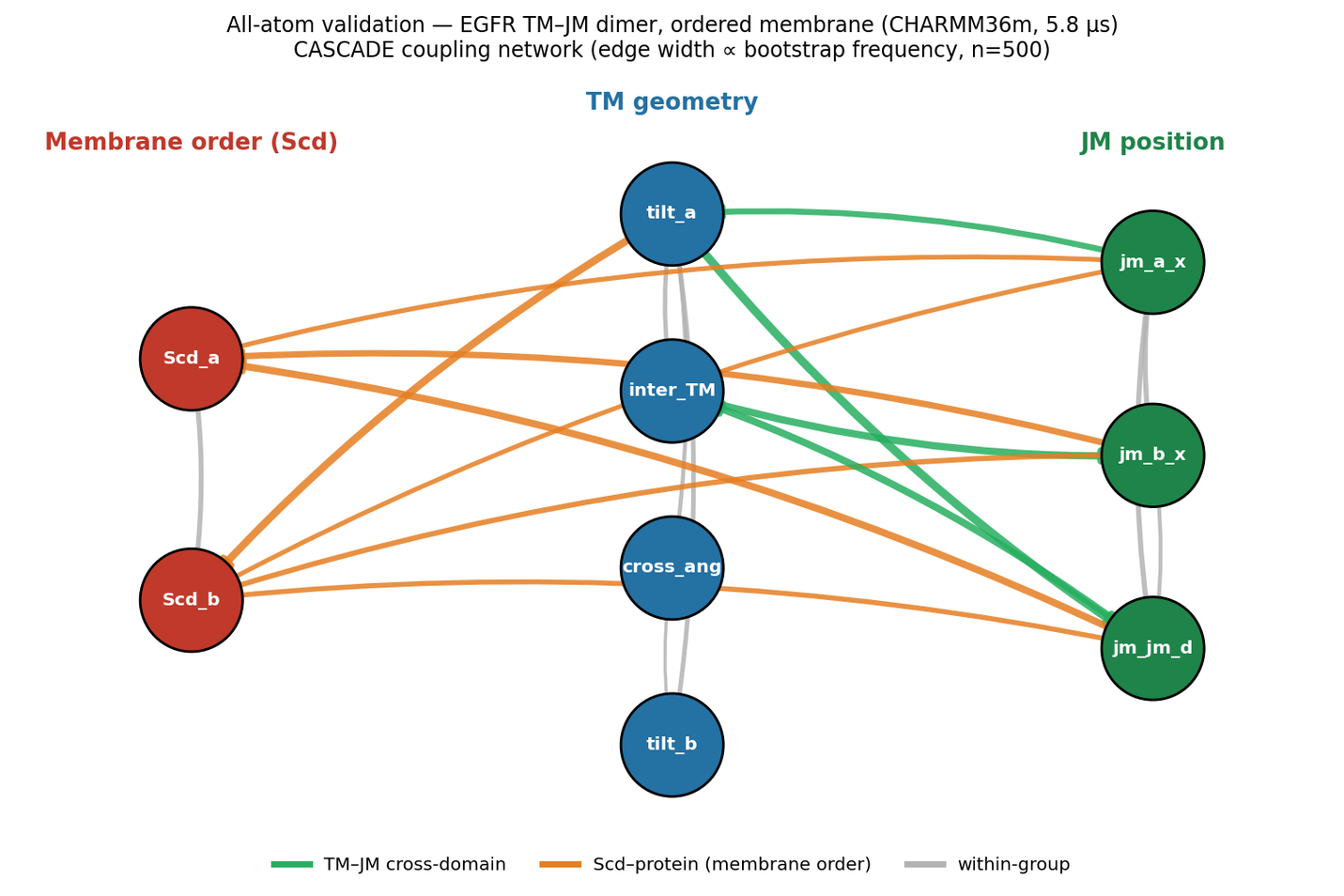

*Figure S3 | All-atom validation: CASCADE coupling network for the EGFR TM–JM dimer in the ordered membrane (CHARMM36m, 5.8 μs; bootstrap n = 500). CASCADE reproduces the same membrane-order hub and TM–JM cross-coupling as the coarse-grained model, with local Scd coupled to four of the seven protein variables (three JM and one TM; bootstrap frequency 0.52–0.91); this validates the coupling detection only (PERI direction is not computed on the all-atom trajectory).*

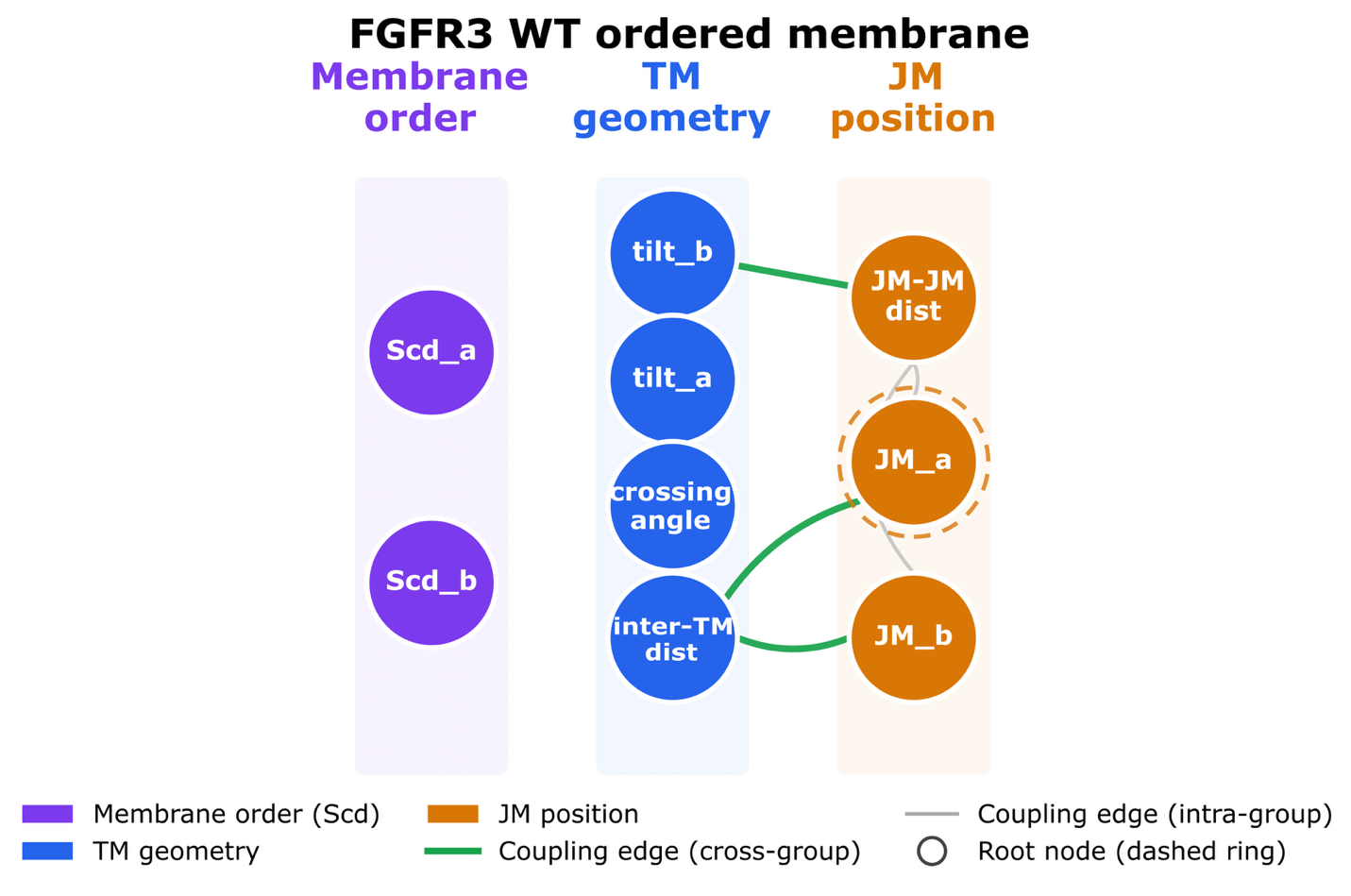

*Figure S4 | CASCADE coupling network for wild-type FGFR3 in the ordered membrane. Internal Bayesian-network skeleton after hierarchical removal of the Scd variance (nodes = structural variables grouped as membrane order, TM geometry, JM position; green = cross-group edges; dashed ring = root node).*

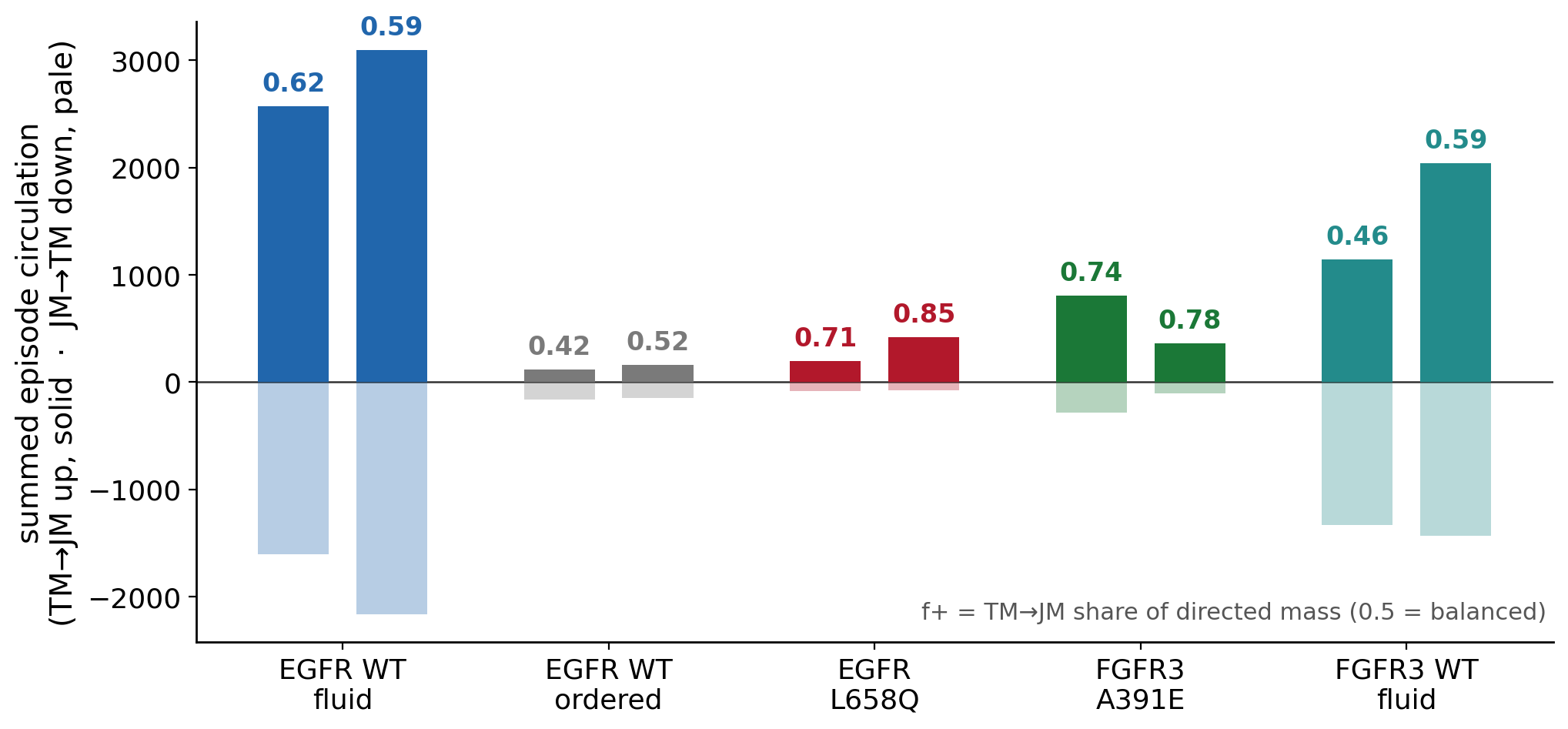

*Figure S5 | Per-trajectory directed-mass balance (0.1 ns), for all ten EGFR and FGFR3 trajectories. For each independent trajectory, the summed circulation of the TM→JM (positive, solid) and JM→TM (negative, pale) episodes; f+ labels the fraction of the total directed mass that is TM→JM. The activating mutants (EGFR L658Q, FGFR3 A391E) and the EGFR fluid wild type are biased toward TM→JM; the ligand-dependent ordered wild type and wild-type FGFR3 in fluid are balanced, their two trajectories falling on opposite sides of 0.5.*

### **Supplementary Note S3. Temporal resolution, aliasing, and statistical inference**

#### **S3.1 The directed coupling is scale-dependent; the fast component is aliased at 1 ns**

The circulation in the (TM, JM) plane is not single-scale. Decomposing the per-episode circulation of one EGFR wild-type fluid trajectory into frequency bands, by bandpass filtering in the Fourier domain and then taking the time-domain signed area of each band (Figure 4a), resolves two directed components of opposite sense: a fast rotation above ≈0.5 ns⁻¹ that is TM→JM (band contribution +915) and dominates the full-bandwidth total of +968, and a slow rotation below ≈0.5 ns⁻¹ that is JM→TM (band contribution −217); because the circulation is a bilinear functional, these band contributions include cross terms and do not sum exactly to the full-bandwidth total. A sampling interval Δt resolves frequencies only up to the Nyquist limit,

*f_Ny = 1 / (2 Δt),*

so at Δt = 1 ns, f_Ny = 0.5 ns⁻¹: the fast TM→JM rotation is undersampled, its apparent frequency folds to |f_rot − m/Δt| (a stroboscopic wagon-wheel alias) and its sign inverts, leaving the slow JM→TM rotation to dominate the apparent direction. At Δt = 0.1 ns, f_Ny = 5 ns⁻¹ resolves the fast rotation. On identical episodes, changing only the sampling interval:

*Table S5 | Sampling-interval dependence: the same episodes read TM→JM at 0.1 ns and JM→TM at 1 ns (aliasing of the fast sub-nanosecond rotation).*

| **Sampling of the same episodes** | **pooled net** | **direction** |
| --- | --- | --- |
| 0.1 ns (full) | +968 | TM→JM |
| 1 ns (every 10th frame) | −332 | JM→TM |

The two are of opposite sign, not a loss of power but the two timescales being read out differently: 1 ns samples the slow JM→TM rotation (and aliases the fast one), whereas 0.1 ns resolves the fast TM→JM rotation that dominates the full-bandwidth net. Because the fast rotation is the direct intramolecular coupling between the two covalently connected ends of one peptide, the TM–JM relationship this study measures, all directivity results use 0.1 ns geometry; the slow JM→TM component is real but a distinct, slower process (Note S3.4 discussion; main-text Discussion).

*The frequency decomposition (a) and cumulative-circulation transient (b) referenced here are shown as Figure 4 in the main text.*

#### **S3.2 Both timescales are real; 0.1 ns is the finest available sampling**

The 1 ns result is therefore not an artifact to be discarded but the genuine slow (multi-nanosecond) JM→TM circulation; the 0.1 ns result is the genuine fast (sub-nanosecond) TM→JM circulation that dominates the net. The direction measured depends on the timescale probed. Every statistic computed on the 1 ns samples returns JM→TM (lag-1 circulation; lagged cross-correlation asymmetry at lags {1}, {2}, {3}, {2–5}, {5,8,12} ns), consistent with those samples reading the slow component. Because 100 ps is the trajectory save interval, 0.1 ns is the finest available sampling and the fast rotation cannot be resolved further; the frequency decomposition (Figure 4a), not a convergence test, is what shows the fast TM→JM component lies above the 1 ns Nyquist limit.

#### **S3.3 Direction does not require the membrane regression**

Because the circulation is built from one-step increments and the membrane-order signal is slow, removing it does not change the increments and therefore does not change the direction. On the 0.1 ns EGFR wild-type fluid trajectory the circulation is identical on the membrane-regressed residuals and on the raw geometry (pooled net = +968 in both). Consequently the 0.1 ns extraction computes only the geometric observables, with local membrane order neither computed nor regressed, a substantial reduction in cost that makes 0.1 ns extraction tractable.

#### **S3.4 Hierarchical Bayesian inference: direction as a posterior probability**

To state the result as direct probability statements, we placed a two-level Bayesian bootstrap on the episode circulations: within each trajectory the episodes receive Dirichlet(1) weights, and the independent trajectories of a condition receive Dirichlet(1) weights (the trajectory being the unit of replication), and the directed-mass fraction f+ is recomputed for each posterior draw (40,000 draws). The procedure is nonparametric: no prior is placed on the effect itself, only the noninformative Dirichlet weights on the observed units. It yields the posterior distribution of f+ for each condition and of any contrast, so that direction is expressed as P(f+ > 0.5) = P(TM→JM) and the mutation effect as P(f+ of the activating mutants − f+ of the ordered wild type > 0).

*Table S6 | Hierarchical Bayesian condition-level posteriors (two-level Bayesian bootstrap): posterior mean f+, 95% credible interval, and P(TM→JM).*

| **Condition** | **posterior mean f+** | **95% credible interval** | **P(f+ > 0.5) = P(TM→JM)** |
| --- | --- | --- | --- |
| EGFR fluid WT | 0.60 | 0.46 – 0.73 | 0.93 |
| EGFR L658Q | 0.76 | 0.42 – 0.94 | 0.95 |
| FGFR3 A391E | 0.75 | 0.54 – 0.89 | 0.99 |
| activating mutants (pooled) | 0.75 | 0.55 – 0.89 | 0.99 |
| EGFR ordered WT | 0.48 | 0.22 – 0.77 | 0.45 |
| FGFR3 fluid WT | 0.53 | 0.39 – 0.67 | 0.65 |

The activating mutants are TM→JM with posterior probability 0.95 (L658Q) and 0.99 (A391E); the ordered wild type is balanced (P = 0.45). For the central contrast, the posterior probability that an activating TM mutation increases the TM→JM bias over the ordered wild type is P(Δf+ > 0) = 0.94 (Figure 5b). These posterior probabilities are the primary inference; the per-trajectory frequentist bootstrap is underpowered (each trajectory yields few episodes) and often does not reach significance on its own, so it is reported only as a consistency check alongside the trajectory-level sign agreement (Notes S2.4–S2.6; Results). Being probability statements about the effect itself rather than about data under a null, they are the natural summary under the small-sample, heterogeneous conditions of membrane-protein simulation.

### **Supplementary Note S4. Robustness controls**

#### **S4.1 The measured direction is not imposed by the simulation restraints**

structure restraint network. The only restraints acting on the protein are the standard MARTINI 2.2 secondary-structure-dependent backbone bonded terms (helical backbone bonds and angles, and i→i+3 backbone dihedrals); these are strictly intrachain and local. Three properties of the topology make the argument concrete. First, every variable that carries the directional signal (the four TM-geometry variables, namely the two helix tilts referenced to the membrane normal, the crossing angle and the inter-TM distance, and the three JM-position variables jm_a_x, jm_b_x and jm_jm_distance) is an inter-chain or protein–membrane quantity that no local, intrachain bonded term in the topology constrains. Second, the backbone dihedral restraints are interrupted at the TM–JM hinge, so the very degree of freedom whose coupling the analysis measures carries no restraint. Third, the restraint scheme is identical in the fluid and the ordered membrane and in wild type and mutant; a factor invariant across a contrast cannot generate the contrast. Independently, (i) the all-atom CHARMM36m trajectory, which carries no secondary-structure restraint of any kind, reproduces the same CASCADE coupling (PERI direction is not applied to the all-atom trajectory, because the dimer remains continuously associated (a single episode) and the episode-based analysis requires discrete association events; it validates the coupling detection only), and (ii) on synthetic AR residuals with matched autocorrelation but no coupling, the circulation returns a null direction at its calibrated false-positive rate (Note S2.5).

#### **S4.2 Stationarity**

Association episodes are the analysis windows; each is a contiguous bound period. Episodes were screened for gross non-stationarity in the individual structural variables (augmented Dickey–Fuller and Spearman monotonic-trend tests) and windows with clear structural drift were excluded. This screen tests each variable for a drifting mean; it does not, and cannot, remove the directed rotational bias, which is a property of the two-variable dynamics rather than a trend in either variable alone. Consistently, the circulation accumulates early within an episode and then saturates (Figure 4b): the cumulative Σc(t) rises well above the constant-rate (diagonal) line, reaching most of its final value within the first fifth of the episode. This front-loaded, saturating accumulation is the signature of a transient directed relaxation (the dimer carries a directional bias while it is still settling, and the bias decays as it settles), not of a constant-rate stationary current. A nonzero stationary current would require sustained external drive (a non-equilibrium steady state), which is absent in these unbiased, ligand-free simulations. The observation of a transient (not steady) circulation is exactly what the non-equilibrium-relaxation interpretation of Note S2.2 predicts, and it resolves the apparent tension between that interpretation and the stationarity screen.

#### **S4.3 A single dimer pair per analysis set**

Dimer detection is part of the episode-extraction step, which is performed at 0.1 ns: all protein pairs are scanned, a dimer requires the TM center-of-mass distance below a 20 Å cutoff, and the single longest continuous dimer event is selected, so an analysis set is never a mixture of two dimer configurations. Each 0.1 ns episode file therefore carries the one pair it represents; for the EGFR ordered condition both trajectories contain a single pair (PROA–PROD, 11 episodes; PROA–PROC, 10 episodes). This is confirmed independently by the all-pairs TM center-of-mass distances on the ordered1 trajectory: only PROA–PROD falls below the 20 Å cutoff (median 12 Å; a dimer in 97% of frames), while the other five pairs never approach it (minimum separations 46–134 Å). The balanced ordered result is therefore that of a single, well-defined dimer, not an average over two oppositely directed dimers. As a further cross-check, the hierarchical decomposition run separately at 1 ns splits its output by pair only when two simultaneous dimers form: it produced a single set for each EGFR ordered trajectory, and pair-specific sets only for the two systems that did form two dimers (FGFR3 wild-type ordered, into AC and BD; FGFR3 A391E, into AD and BC), each analyzed as an independent dimer. For FGFR3 A391E the AD pair is the dominant, well-formed dimer (16 episodes at 0.1 ns), whereas the BC pair associated only transiently (two episodes even at 1 ns) and is too sparsely sampled to support a direction call; the reported A391E result is therefore the well-populated AD dimer, not a selection among comparable pairs. FGFR3 wild-type ordered was excluded from the quantitative comparison for independent reasons (near-continuous association yielding few discrete episodes). The per-condition trajectories used are those listed in Table S3.

### **Supplementary Note S5. Molecular dynamics simulation details**

Coarse-grained simulations: GROMACS 2023.3, MARTINI 2.2 force field, MARTINI polarizable water, 0.15 M NaCl, 20 fs timestep, velocity-rescale thermostat (298 K, τ_T = 1.0 ps; protein, membrane and solvent coupled separately), semi-isotropic Parrinello–Rahman barostat (1 bar, τ_P = 12 ps, compressibility 3 × 10⁻⁴ bar⁻¹), reaction-field electrostatics (relative permittivity 2.5, Coulomb cutoff 1.1 nm), van der Waals cutoff 1.1 nm with potential-shift modifier; Verlet scheme, neighbour list every 20 steps. Inputs generated with CHARMM-GUI Martini Maker, standard production parameters unchanged. Protein converted with martinize (v2.5), martini22p parameter set, DSSP-assigned secondary structure. No elastic network is applied: the only protein restraints are the standard MARTINI 2.2 secondary-structure-dependent backbone bonded terms (helical bonds and angles; i→i+3 dihedrals, φ₀ = −120°, k = 400 kJ mol⁻¹). No distance restraint is derived from a reference structure; the dihedral restraints are interrupted at the TM–JM hinge and no term acts on inter-chain geometry. CHARMM-GUI POSRES restraints appear only inside an #ifdef block and are inactive in production. EGFR TM–JM: residues 634–683 (TM helix 646–668; intracellular JM 669–683; PDB 2M20). FGFR3 TM–JM: corresponding segment (PDB 2LZL). Mutations via CHARMM-GUI. Fluid membrane: DIPC(70)/DOPS(30), 8×8×15 nm³, 2 peptide copies, 10 μs. Ordered membrane: upper leaflet DIPC/DPSM/CHOL equimolar, lower leaflet DIPC/CHOL/DOPS equimolar, 40×40×15 nm³, 4 peptide copies, 10 μs. Each system reached 10 μs by successive extensions with identical parameters. All-atom: CG snapshot at 10 μs backmapped to CHARMM36m (no secondary-structure restraint of any kind), 10×10×10 nm³, 5.8 μs. Trajectories saved every 100 ps; geometric observables for the direction analysis extracted at 0.1 ns. Dimers: TM backbone minimum distance < 6 Å sustained ≥ 100 ns (association episodes). Quality control: area per lipid (fluid 0.64, ordered 0.51 nm²), thickness (fluid 3.8, ordered 4.2 nm), DPSM Scd 0.35–0.45 (Lo confirmed), backbone RMSD 0.3–0.5 nm after 100 ns equilibration.

Software: GROMACS 2023.3, MARTINI 2.2, Python 3.10, MDAnalysis 2.7, PyMC 5.10, NumPy 1.26, SciPy 1.12, statsmodels 0.14. Circulation and synthetic-validation code will be deposited on acceptance.
